# Insulators As Dynamic, Tunable Regulators of Enhancer-Promoter Coordination in Living *Drosophila* Embryos

**DOI:** 10.64898/2026.09.09.750400

**Authors:** Noel Buitrago, Cesar Nieto, Abhyudai Singh, Bomyi Lim

## Abstract

The three-dimensional organization of the genome enables enhancers and promoters to interact across vast distances and direct transcription. Yet whether architectural elements, such as insulators, serve as rigid, passive barriers or as dynamic, active organizers of this communication remains unclear. Here, using single-cell, live imaging of a *Drosophila* transgene in which a single enhancer regulates two equidistant promoters, we confirm that the enhancer engages both promoters simultaneously and show that coordinated bursting is intrinsically more productive than uncoordinated activity. Flanking this system with insulators increases coordinated bursting frequency and transcriptional output, indicating that insulator-mediated looping promotes multi-way enhancer-promoter interaction. Further, bidirectionally-paired, homotypic insulators produce stronger coordination than unidirectional pairs. Inserting an intermediate insulator to generate competing loop configurations, together with two-state promoter modeling, we show that these chromatin loops are highly dynamic. This work reframes insulators as active, tunable regulators that shape the frequency, coordination, and productivity of enhancer-promoter communication.

## Introduction

Enhancer regions are characteristic elements of the metazoan genome, responsible for the spatiotemporal regulation of gene expression. These non-coding, *cis*-regulatory sequences contain binding sites for transcription factors (TFs) that can either augment or repress transcription, and interact with the promoter of their target gene to carry out their function.^1^ The exact mechanism by which the enhancer is able to communicate with its target gene has been an ongoing area of research, and yet no clear spatial or temporal thresholds between the enhancer-promoter (E-P) interactions that are required to initiate transcription have been elucidated. Several models of E-P communication have been proposed, including loop extrusion and phase separation, but no consensus has emerged.^2,3^ In *Drosophila* embryos, live imaging revealed that transcription initiated only when the enhancer and promoter came within ∼400 nm of each other;^4,5^ however, E-P distance in mouse cells was not necessarily closer at actively transcribing loci compared to inactive ones.^6,7^

In practice, robust gene expression rarely results from such isolated, one-to-one E-P pairs; instead, it typically involves complex multi-way interactions, wherein a single gene may be regulated by numerous enhancers or a single enhancer may regulate multiple target genes.^1^ Such examples are the *even-skipped* (*eve*) locus in *Drosophila* where five enhancers direct expression of the *eve* gene into a seven stripe pattern^8^ or the β-globin locus in mammals where a single upstream Locus Control Region (LCR) enhancer directs expression of adult hemoglobin gene clusters.^9^

Of these two regulatory configurations (multiple enhancers converging on a single gene versus a single enhancer coordinating multiple genes), the mechanism of the latter remains comparatively less understood. Nevertheless, this arrangement is far from rare: in *Drosophila* alone, about 50 pairs of paralogous genes are thought to share regulation by a common enhancer.^10^ Live-imaging studies have provided a great deal of insight into how a single enhancer can coordinate multiple target genes. Distal paralogs such as *scylla* and *charybde* are positioned 235 kb apart in the fruit fly genome but nonetheless share a regulatory element that spatiotemporally coordinates their transcriptional activity.^10^ Shorter-range, transgenic studies have also revealed a high level of coordinated gene activity when two promoters are positioned near to a single enhancer.^11^ Such results present strong evidence that the single enhancer interacts with multiple promoters to initiate simultaneous transcriptional activation, i.e. co-activation. However, coordinated activity could also arise independently of direct enhancer action: for instance, through stochastic transcriptional bursting that aligns by chance, or through global regulatory factors (e.g., shared transcriptional machinery or cell-cycle stage) that influence many genes simultaneously. Consequently, the question arises of how this multi-way E-P interaction is achieved and with sufficient frequency to ensure proper development and cellular maintenance.

Regulatory complexity is compounded by the fact that enhancers and the genes they regulate are tens, even hundreds, of kilobases apart in linear genomic distance.^12^ Far from empty space, these intermediate regions are riddled with other regulatory elements and non-target promoters. While the precise mechanisms governing E-P specificity and frequency remain unresolved, it is well established that chromatin is not organized randomly in the nucleus. Rather, its three-dimensional architecture is highly regulated, such that this organization specifically facilitates appropriate E-P interactions, while precluding undesirable ones.^10,13^

Essential to establishing this organization are insulator elements. Insulators are intergenic sequences that lack inherent enhancer activity, but instead contain binding sites for specific insulator proteins. Some elements, such as Homie/Nhomie in *Drosophila*, directly pair with one another to form loop-like domains that drive local chromatin topology.^14–16^ Other insulators, such as CCCTC-binding factor (CTCF) in vertebrates, serve as endpoints for loop-extruding protein complexes.^17^ Regardless of mechanistic differences, insulators have long been theorized to organize chromatin into ‘higher order domains’.^18^ The advent of chromatin conformation capture technologies (e.g. 4C, Hi-C, Micro-C) has since validated their function, implicating insulators specifically at the boundaries of topologically associating domains (TADs).^13^ These domains represent regions of the genome with high internal contact frequency. The boundary elements, or insulators, delimiting TADs both facilitate intra-TAD interactions and inhibit inter-TAD interactions, a property demonstrated in numerous classical enhancer-blocking assays.^19–21^ By constraining which enhancers and promoters share a topological neighborhood, TAD boundaries therefore directly influence which regulatory interactions are permitted to occur. Thus, insulators represent a crucial class of architectural elements, responsible for the partitioning of the genome into distinct ‘neighborhoods’ of transcriptional activity.

Despite these advancements in characterizing insulator function at topological boundaries, a knowledge gap remains regarding their individual contributions in modulating multi-way E-P interactions and subsequent transcription. Current research largely focuses on endogenous loci with considerable variation in insulator position, orientation, density and type. Such complexity muddies the contributions of a single insulator element or insulator pair. It is still unclear whether insulators form rigid, physical barriers that block regulatory cross-talk or if their properties can be optimized to fine-tune the frequency and stability of multi-way E-P interactions. Moreover, it remains unresolved whether insulators act permissively, enabling qualitatively new E-P contacts that would not otherwise occur, or whether they instead modulate the kinetics of interactions that are already accessible to the enhancer.

To bridge these gaps in understanding, we employed a highly-controlled transgenic system in living *Drosophila* embryos consisting of a single enhancer regulating two equidistant target genes, into which we systematically introduced insulator elements in varying positions and orientations. Using the MS2/MS2 Coat Protein (MCP) and PP7/PP7 Coat Protein (PCP) systems to monitor the transcriptional activity of individual nuclei in real time, we find that flanking insulators increase the degree of coordinated transcriptional bursting and total mRNA output, with coordinated bursts proving intrinsically more productive than uncoordinated ones. Both of these effects were strongly dependent on the relative orientation of flanking insulators, with homotypic insulators preferentially pairing when oriented opposite of one another. Introducing an intermediate insulator between the enhancer and one target promoter further revealed that this looping is highly dynamic, with the system adopting multiple topological configurations within a single nuclear cycle, consistent with proximity-based insulator pairing. Spatial isolation of the PP7-tagged promoter also bolstered the transcriptional output of the neighboring MS2-tagged promoter, revealing an opportunistic relationship between them. Together, these findings reframe chromatin organization not as a static scaffold but as a dynamic, tunable layer of transcriptional regulation whose topology and productivity are inextricably linked.

## Results

### A single enhancer can activate transcription of two equidistant genes simultaneously

To investigate the nature of multi-way E-P interaction in a controlled environment, we employed a previously reported transgenic construct, hereafter referred to as ‘Dual’.^11^ This system consists of a single *snail* distal enhancer (*sna*DE) positioned in *cis*, downstream of two equidistant *snail* promoters (*sna*Prs), each driving transcription of a full *yellow* reporter gene (Figure 1A). In *Drosophila*, the *snail* (*sna)* gene is a critical determinant of mesoderm specification and whose distal enhancer drives robust, uniform transcriptional activity across the ventral mesoderm (Figure S1).^22^ This widespread activation provides a large population of nuclei under equivalent enhancer influence, offering a practical advantage for single-cell live imaging analysis. The sequence between each promoter and the enhancer is nearly identical with two exceptions: 1) the 5’UTR of one reporter contains 24 copies of the *MS2* stem loop sequence and the other contains 24 copies of the PP7 stem loop sequence to distinguish transcriptional activity from each promoter and 2) the insertion of a unique 1kb *lacZ* segment within the intron of each *yellow* reporter gene (Figure 1A). As transcribed, the MS2 and PP7 stem loop structures bind their respective coat proteins, MCP and PCP, with high affinity. By fusing these coat proteins to fluorescent proteins and maternally loading them, we visualize their localization, and thus nascent transcription, in real time within each nucleus.^23–25^ This transgenic construct was integrated into a specific region on the third chromosome (3L) of the *Drosophila melanogaster* genome that is transcriptionally silent and free of confounding regulatory architecture.^26–28^ In doing so, the observed behaviors could be largely attributed to that of the system as opposed to the surrounding genomic context. Live imaging occurred for the entirety of nuclear cycle 14 (nc 14), a specific developmental window 2-3 hours after fertilization and demarcated by the end of the thirteenth mitosis and the onset of gastrulation.^24,29^

**Figure 1.**
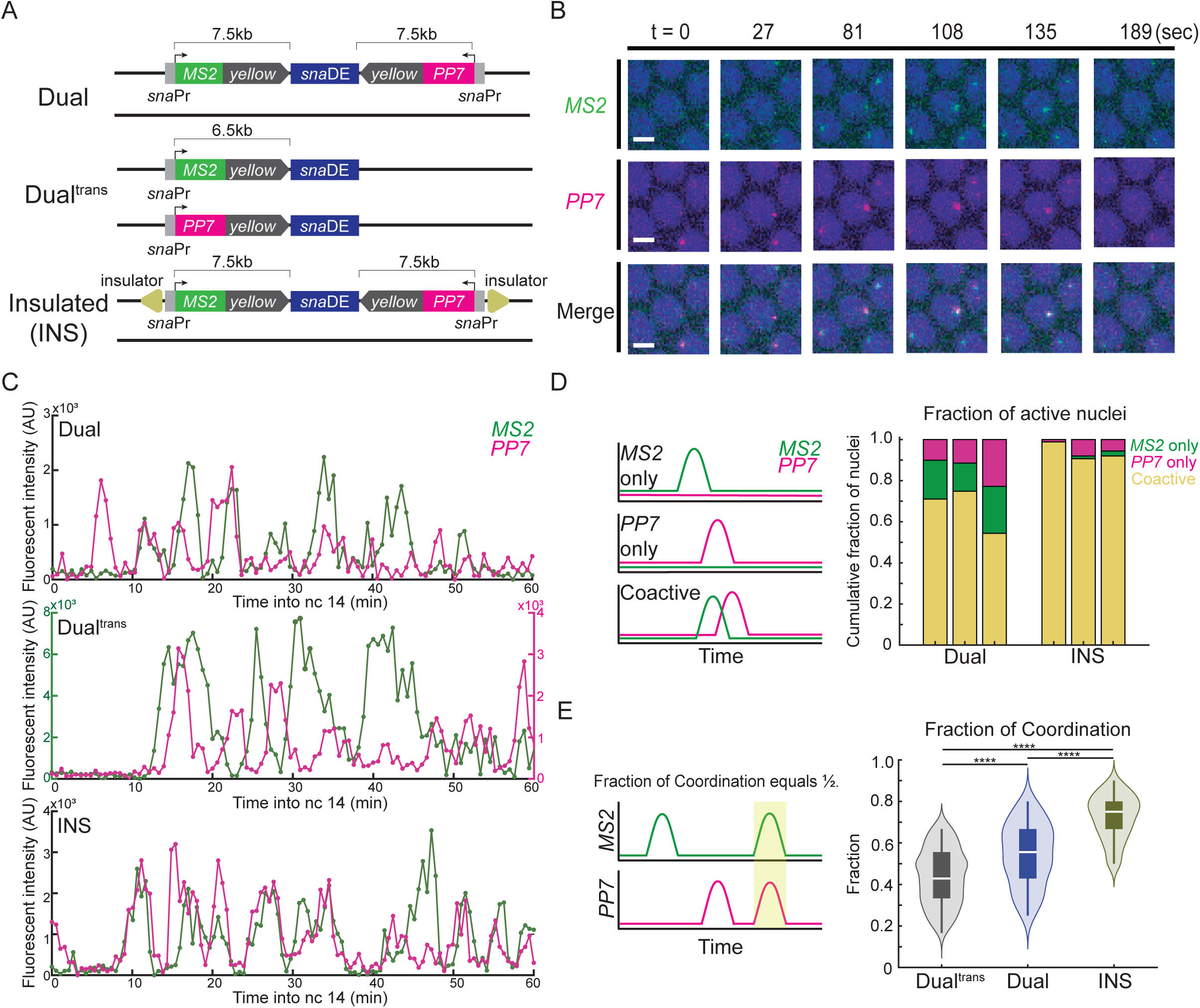
Flanking insulators facilitate E-P interaction and synchronize transcriptional bursting events. (A) Schematic of transgenic constructs: Dual, Dual^trans^, and INS. (B) Live-imaging snapshots showing a coordinated transcriptional bursting event in a single nucleus of the Dual construct over time. Scale bar (white rectangle) represents 5um. (C) Representative MS2 (green) and PP7 (magenta) trajectories for single nuclei across nc 14 from Dual (top), Dual^trans^ (middle) and INS (bottom). (D) Left: Schematic for classification of nuclei as either MS2 only, PP7 only, or coactive. Right: Bar chart displaying cumulative fraction of nuclei that fall within each category (N = 3 biological replicates for each construct, one bar per replicate). INS construct shows greater proportion of coactive nuclei. (E) Left: Illustration of the Fraction of Coordination metric. Example MS2 and PP7 trajectories from a single nucleus show one uncoordinated burst and one coordinated burst highlighted in yellow. The resulting Fraction of Coordination is 0.5. Right: Plot of Fraction of Coordination for Dual^trans^, Dual and INS (N = 3 biological replicates for each construct). Number of nuclei in each data set is as follows: n(Dual^trans^) = 674, n(Dual) = 361, n(INS) = 566. ****p<0.0001 by one-way ANOVA and Tukey-Kramer test (E,right).

As demonstrated previously with this system, both reporters were expressed within the appropriate *sna* expression domain, with MS2 and PP7 signals each displaying discontinuous, bursty transcriptional activity throughout nc 14 (Figures 1B, 1C; Supplemental Movie 1).^11^ Overall, between 50-70% of transcriptionally active nuclei displayed both MS2 and PP7 activity at some point within nc 14, i.e. coactivity (Figure 1D). While this result confirms that both reporter genes have access to the enhancer during our observation window, it lacks the temporal resolution to distinguish between distinct biophysical models of E-P interaction. Given the 45-minute duration of nc 14 relative to the minutes-long transcriptional bursts of the *sna* enhancer, a high proportion of coactive nuclei could plausibly arise from purely sequential E-P engagement. Whether true simultaneous co-activation occurs with sufficient frequency to suggest spatial co-occupancy, rather than rapid sequential contacts, remains unresolved. While coordinated bursting was reported previously in this system, it was quantified per reporter rather than as a unified metric. Further, coordination was assessed through randomization of experimental traces rather than a genuine biological control for independent regulation.^11^

To address these limitations, we established a stringent analytical approach to evaluate co-activation kinetics. We used a unified metric, Fraction of Coordination, defined as the number of coordinated bursting events divided by the total number of bursting events in the less bursty reporter. Our analysis revealed that coordinated bursting is a predominant feature of the system rather than a rare event, as evidenced by a high Fraction of Coordination (>50%) (Figure 1E). To further rule out the possibility that this reflects stochastic bursting rather than true *cis*-mediated co-activation, we decoupled the two reporters onto homologous alleles, each independently regulated by a downstream *sna*DE at a comparable E-P distance (Figure 1A). This arrangement, termed Dual^trans^, maintained its high level of transcriptional activity; however, it demonstrated a significantly lower incidence of coordinated bursting, confirming that the elevated coordination observed in Dual reflects genuine co-regulation by a single shared enhancer, rather than coincidental co-bursting (Figure 1E). Our results show that an enhancer can activate two target promoters at the same time, resulting in coordinated transcriptional bursting. This simultaneous engagement of a single enhancer and two promoters suggests spatial co-occupancy or a transcriptional hub of these regulatory elements, rather than rapid sequential contacts or a ‘kiss and kick’ form of E-P interaction.

### Insulator-mediated looping synchronizes gene activity in an orientation-dependent manner

The high degree of coordinated bursting observed in the Dual construct raised the question of whether local chromatin architecture actively facilitates multi-way E-P interaction. Indeed, the genomic landing site used in these experiments (vk33) was suggested to harbor endogenous insulator activity that may contribute to the looping of the system’s regulatory elements,^11^ and yet chromatin conformation capture techniques revealed no clear topological domains.^27,28^ To directly test whether insulator-mediated looping modulates the degree of multi-way E-P interaction and its transcriptional consequences, we systematically introduced insulator elements in varying positions and orientations into the Dual construct. Specifically, we employed the well-characterized Su(Hw)-dependent *gypsy* insulator, which originates from the 5’UTR of the *gypsy* retrotransposon and displays classical insulator properties including modulation of E-P interaction and barrier activity against heterochromatin spread.^30–32^

We inserted diverging *gypsy* insulators, consisting of 12 tandem copies of the Su(Hw) insulator binding sequences, at the flanking positions of the Dual construct to create the Insulated (INS) construct (Figure 1A). In theory, these elements should pair with one another, bringing all elements of the system within proximity and promoting their interaction. Consistent with this idea, we observed heightened levels of coactivity (90%) in INS compared to Dual (65%), indicating that the enhancer could indeed more reliably find both promoters within nc 14. Thus, insulator-mediated looping of our system appears to have reduced the 3D search space of the enhancer. Moreover, the expression of MS2 and PP7 became further temporally aligned with ∼75% of transcriptional bursting events being coordinated compared to ∼55% without insulators (Figures 1D and 1E). This heightened synchronization suggests that the insulator-mediated spatial proximity also promotes the formation of a single, shared transcriptional hub that both promoters access simultaneously.

Importantly, the degree of coordinated bursting was largely uniform across the *sna* expression domain, with no spatial preference for higher or lower coordination. Furthermore, the domain of expression itself remained comparable in size to that of the uninsulated Dual, indicating that the introduction of insulators did not ectopically expand enhancer activity (Figures S2A and S2B). Therefore, the enhanced coordination reflects an intrinsic property of insulator-mediated chromatin organization, rather than a downstream consequence of spatially varying transcription factor concentrations (e.g., Dorsal morphogen gradients). To test whether insulators alone direct ectopic transcriptional activity independent of the enhancer, we removed the enhancer while retaining the insulators, called INS^Δ*sna*DE^, and observed very little to no transcriptional activity (Figures S2C and S2D), confirming that the coordination effects of INS require the enhancer itself. Together, these findings highlight chromatin organization and enhancers as independent layers of transcriptional regulation, where insulators regulate the likelihood and timing of E-P interactions, but do not affect the enhancer’s intrinsic regulatory capability.

Having established that insulator pairing enhances multi-way E-P coordination, we next asked whether this effect depends on the relative orientation of the flanking insulators. Although *Drosophila* insulators have been increasingly recognized as orientation-dependent chromatin organizers, the *gypsy* element specifically has been characterized as largely orientation-independent in insulator bypass assays;^33^ however, these studies did not assess its functional impact on transcriptional kinetics. To directly assess whether orientation influences homotypic insulator pairing and its downstream transcriptional consequences, we generated two additional constructs with flanking insulators: 1) a convergent arrangement (CONV), which is predominant among CTCF-mediated topological boundaries within vertebrate genomes^34^ and 2) the unidirectional arrangement (UNI), where both insulators are oriented in the forward direction (Figure 2A). Live imaging of these constructs revealed that the high levels of coactivity and coordination were maintained when the insulators were placed in the convergent orientation, but not in the unidirectional configuration (Figures 2B and 2C). The bidirectional insulator arrangements, INS (upstream insulator reverse, downstream forward) and CONV (upstream forward, downstream reverse), displayed superior levels of coactivity and coordination, indicating improved loop formation or stability (Figures 2B-2E). The UNI construct still significantly elevated coordination relative to the uninsulated Dual, although its effect was markedly more modest than that of the bidirectional configurations (INS and CONV) (Figure S2E). As an orthogonal validation of our coordination metric, we also calculated the Spearman’s correlation coefficient between the MS2 and PP7 trajectories of each coactive nucleus across all constructs. This alternative analysis showed a highly consistent trend wherein the transcriptional activity of the two reporters correlated more strongly with flanking insulators present than in the uninsulated Dual, though the unidirectional arrangement again yielded a notably weaker effect (Figures 2F and S2F).

**Figure 2.**
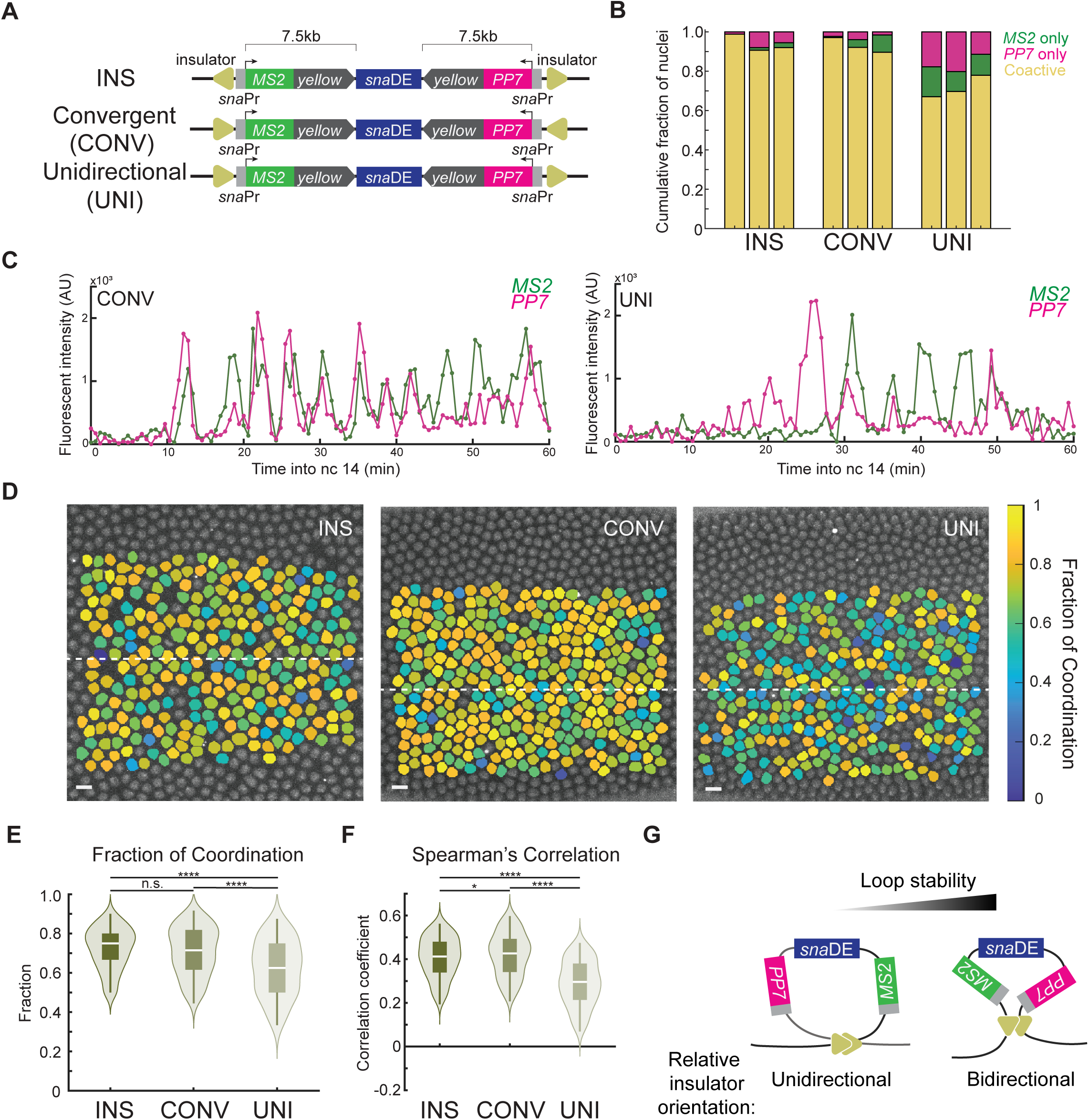
Insulator-mediated coordination of gene activity is dependent on relative orientation. (A) Schematic of transgenic constructs with varying relative orientation of flanking insulators. (B) Bar chart displaying cumulative fraction of nuclei that show only MS2 activity (green), only PP7 activity (magenta) and coactivity (yellow). INS and CONV show greater proportions of coactive nuclei compared to UNI. (C) Representative MS2 (green) and PP7 (magenta) trajectories from CONV (left) and UNI (right). (D) False-colored snapshot from imaging depicting Fraction of Coordination of each coactive nucleus in the pictured embryo. White dashed lines represent the ventral midline of the embryo. There is no apparent difference in the width of the domain nor any spatial arrangement for nuclei with higher or lower coordination. Scale bar (white rectangle) represents 5um. (E,F) Violin plot of (E) Fraction of Coordination and (F) Spearman’s correlation coefficient between the MS2 and PP7 trajectories of individual coactive nuclei (N = 3 biological replicates for each construct). Number of nuclei in each dataset is: n(INS) = 566, n(CONV) = 674, n(UNI) = 507. (G) Illustration of chromatin looping conformations and their stability. Unidirectional homotypic insulator pairing yields circle loop structure whereas bidirectional pairing produces stem loop structure. The stem loop appears more stable than the circle loop. n.s. not significant, *p<0.05, **** p<0.0001 by one-way ANOVA and Tukey-Kramer test (E and F).

Together, these data demonstrate that homotypic *gypsy* insulator pairing in *cis* is orientation-dependent, with bidirectional arrangements (INS and CONV) producing significantly stronger synchronization of transcriptional bursting than unidirectional ones (UNI). While all insulator configurations elevated coordination above the uninsulated baseline, the degree of effect varied substantially with orientation, suggesting that relative insulator orientation is a tunable parameter of chromatin loop-mediated gene regulation (Figure 2G).

### Insulators enhance transcriptional output through increased burst frequency and size

Having established that insulator-mediated looping increases the degree of coordinated transcriptional bursting, we next asked whether this heightened coordination has any consequence for transcriptional output. Visualizing the MS2 transcriptional trajectories of all active nuclei as heatmaps, we observed higher fluorescent intensities in the INS and CONV configurations compared to the Dual and UNI constructs (Figure 3A). Quantification of total RNA production by integrating beneath the MS2 and PP7 trajectories confirmed that the bidirectional INS and CONV had significantly greater MS2 and PP7 outputs compared to the uninsulated Dual (Figures 3B and 3C). Notably, this pattern mirrored the trend in coordinated bursting, such that constructs with higher coordination showed greater output (Figures 2E, 2F, 3B and 3C). On the other hand, UNI (unidirectional insulator positioning), which showed only a modest degree of coordination, showed no significant change in MS2 output and only a slight increase in PP7 output (Figures 3B and 3C). The lack of a corresponding output increase in UNI, despite its elevated coordination, suggests that coordination alone may not be sufficient to drive greater transcriptional output. We next asked whether coordination and output were similarly linked at the level of individual nuclei. Regression analysis between the coordination (Spearman’s correlation coefficient) and total output from individual nuclei revealed little to no relationship (Figures S3A and S3B). Thus, a nucleus’s degree of coordination does not scale with its output, despite coordination predicting output differences between constructs. Instead, the enhanced output in INS and CONV likely reflects additional, construct-level changes in transcriptional bursting.

**Figure 3.**
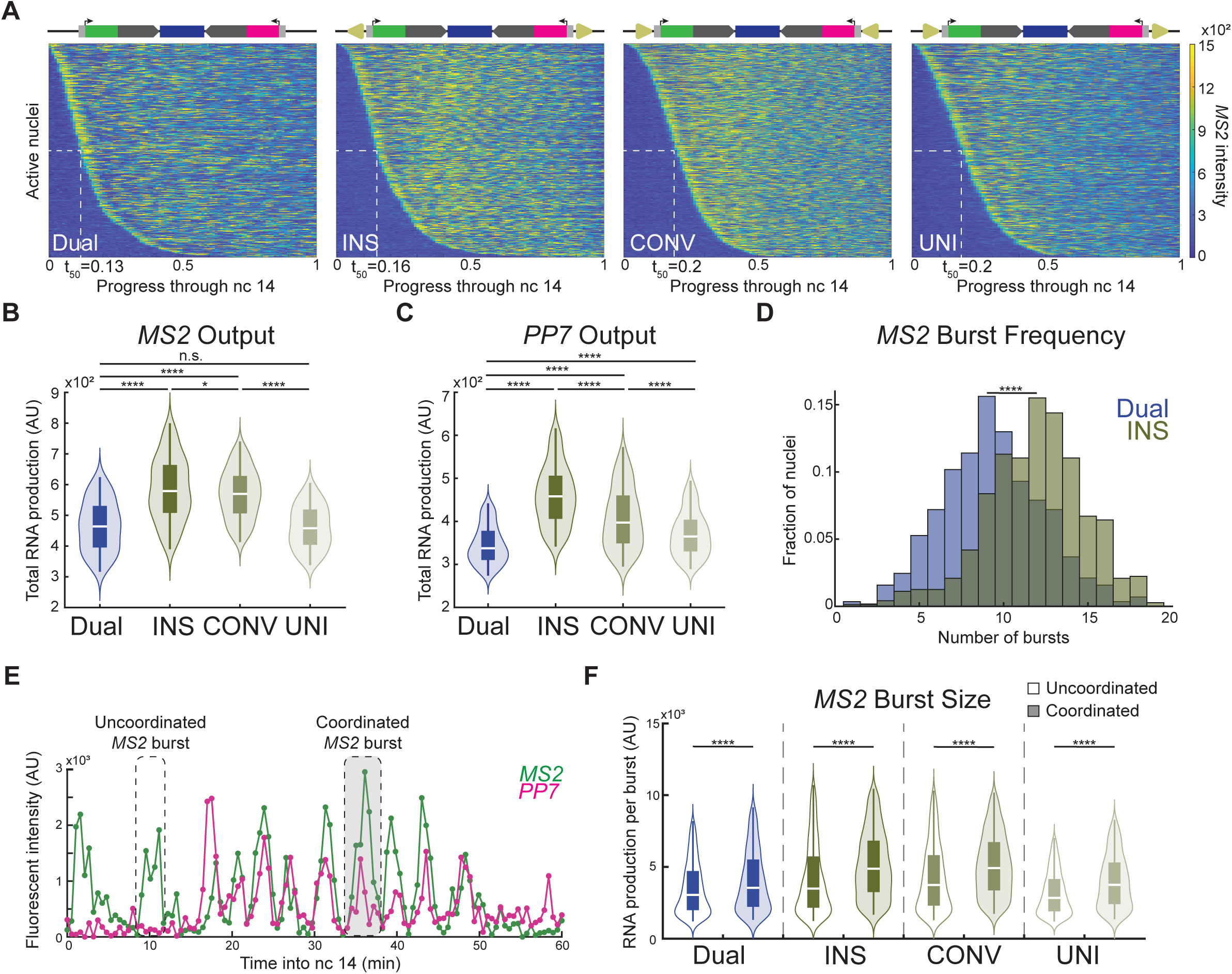
Insulator-mediated looping enhances transcription through burst frequency and size. (A) Heat map of transcriptional trajectories for all active nuclei in each construct sorted by transcriptional onset (N = 3 biological replicates for each construct). Each row represents a single trajectory from an individual nucleus. The x-axis represents normalized progress through nc 14, from 0 (start) to 1 (end). Color scale corresponds to the fluorescent intensity of MS2. The timing of activation of half the total active nuclei is reported as t_50_. (B) Violin plot of total MS2 mRNA produced by individual MS2-active nuclei throughout nc 14 (N = 3 biological replicates for each construct). Number of nuclei in each data set is as follows: n(Dual) = 457, n(INS) = 575, n(CONV) = 702, n(UNI) = 587. (C) Violin plot of total PP7 mRNA produced by individual PP7-active nuclei throughout nc 14 (N = 3 biological replicates). Number of nuclei in each dataset is as follows: n(Dual) = 434, n(INS) = 600, n(CONV) = 692, n(UNI) = 617. (D) Histogram showing the number of MS2 bursts per individual MS2-active nucleus (N = 3 biological replicates for each construct). Plotted with a relative probability normalization such that the height of the bars corresponds to the fraction of nuclei with that number of MS2 burst events during nc 14. Addition of flanking insulators shifts the histogram to the right, indicating an increase in overall burst frequency. The total number of nuclei is as follows: n(Dual) = 570, n(INS) = 760. (E) Representative MS2 (green) and PP7 (magenta) trajectories. Empty callout box shows an uncoordinated MS2 burst. Filled callout box shows a coordinated MS2 burst. (F) Violin plot of MS2 burst size for each construct sorted into uncoordinated (unfilled, left) and coordinated (filled, right) bursts. Burst size data was compiled and sorted from three biological replicates. The number of data points for each set is as follows: n(Dual,Uncoordinated) = 3241, n(Dual,Coordinated) = 1700, n(INS,Uncoordinated) = 2723, n(INS,Coordinated) = 4384, n(CONV,Uncoordinated) = 3698, n(CONV,Coordinated) = 4294, n(UNI,Uncoordinated) = 3948, n(UNI,Coordinated) = 2507. n.s no significant difference, *p<0.05, ****p<0.0001 by one-way ANOVA and Tukey-Kramer test (B,C) or by Wilcoxon rank-sum test (D,F).

We therefore examined the burst frequency and burst size of active nuclei. Burst frequency reflects the rate of transcriptional initiation and is influenced by the frequency of productive E-P engagement, whereas burst size is associated with the rate of RNA Pol II pause-release during the active promoter state.^35^ By quantifying the number of bursts per nucleus during nc 14, we found that the addition of flanking insulators significantly increased the burst frequency of both MS2 and PP7 by 33% and 57%, respectively (Figures 3D, S3C and S3D). This result reinforces the notion that insulator-mediated looping reduces the search space through which the enhancer must explore to find either promoter, hence driving more frequent E-P interaction.^11,28^ We also observed a significantly greater median MS2 and PP7 burst size in the bidirectional INS and CONV configurations compared to the uninsulated Dual (Figures S3E and S3F). Both MS2 and PP7 showed greater burst size despite the presence of two promoters sharing the same enhancer, a scenario in which classical models of promoter competition would predict reduced, not enhanced, output per promoter. This finding suggests that insulator-mediated looping’s effect on transcription is stabilizing rather than merely permissive. The concomitant increase in burst size was unexpected and may reflect a positive feedback mechanism, whereby more frequent bursting promotes greater transcription factor clustering, which in turn stabilizes and prolongs each burst.^36^ Alternatively, stable chromatin looping may stabilize the active promoter state, enhancing Pol II pause-release efficiency and hence increasing burst size independently of contact frequency.

Given that insulators both increased the number of bursts and the proportion that were coordinated, we wondered if coordinated bursts themselves were intrinsically more transcriptionally productive, and if so, whether this required insulator-mediated looping. To investigate this idea, we categorized individual transcriptional bursting events as either ‘Uncoordinated’ or ‘Coordinated’ (Figure 3E). Upon sorting, we observed that coordinated bursts were significantly more sizable than uncoordinated ones (Figure 3F). Interestingly, this trend held even in the case of the uninsulated Dual construct, indicating that this phenomenon is a feature of multi-way E-P interaction rather than a direct effect of local chromatin architecture. Previous work has validated that simultaneous multi-way bursting is preceded by a larger clustering of TFs compared to one-on-one E-P interaction,^36^ suggesting that the enriched transcriptional microenvironment during co-occupancy drives larger bursts from both promoters simultaneously. Importantly, both promoters benefited equally from this enriched microenvironment, arguing against a winner-takes-all model of promoter competition and instead suggesting a synergistic relationship between co-regulated promoters.

### Insulator-mediated looping is highly-dynamic, sampling multiple competing configurations

Given that flanking insulators formed a loop and changed transcriptional behaviors, we sought to determine the stability of this insulator-mediated looping. To address this, we leveraged our existing INS system and inserted a third, intermediate insulator to enable the formation of multiple competing loop configurations. More specifically, we placed an additional *gypsy* insulator immediately between the *sna*DE and the 3’UTR of the PP7-containing *yellow* reporter gene, referred to as ‘PP7-Blocked’ (Figure 4A). In doing so, we expected the intermediate insulator to pair with either flanking insulator, in both cases sequestering the PP7 promoter from the enhancer. This arrangement was highly reminiscent of classical enhancer-blocking studies and we expected to observe little to no PP7 transcriptional activity.^19–21^ Consistent with a partial blocking effect, we observed a significant portion of nuclei that were MS2-only monoactive, indicating that the presence of an intermediate insulator was blocking the interaction of the enhancer and the PP7’s promoter (Figure 4B; Supplemental Movie 2). PP7 activity was almost never observed in isolation: nearly every PP7-active nucleus was also MS2-active (Figure 4B). Surprisingly, about 60% of all active nuclei were still coactive, suggesting that the majority of nuclei escaped insulator-mediated blocking and still retained PP7 activity in spite of an intermediate insulator (Figure 4B). MS2 and PP7 activity were comparably frequent in the absence of the intermediate insulator (Figure 2B). Therefore, the near-total dependence of PP7 activity on concurrent MS2 activity is unlikely to reflect an intrinsic asymmetry between the two promoters. Instead, this pattern is consistent with a model in which the system assumes one of two broad configurations: a PP7-exclusionary state where the intermediate insulator pairs with either flanking insulator to sequester PP7 and permit only MS2 activity, or a system-wide loop in which both reporter genes remain engaged with the enhancer. We also generated a construct where this intermediate insulator was inverted, hereafter referred to as PP7-Blocked^INV^. Because homotypic insulators preferentially pair head-to-head, the intermediate insulator is predicted to pair favorably with the MS2-flanking insulator in PP7-Blocked and the PP7-flanking insulator in PP7-Blocked^INV^ — both yielding a PP7-exclusionary loop, differing only in which promoter is co-enclosed with the enhancer. Consistent with this, PP7-Blocked and PP7-Blocked^INV^ showed similar levels of coactivity (Figures 4A and 4B).

**Figure 4.**
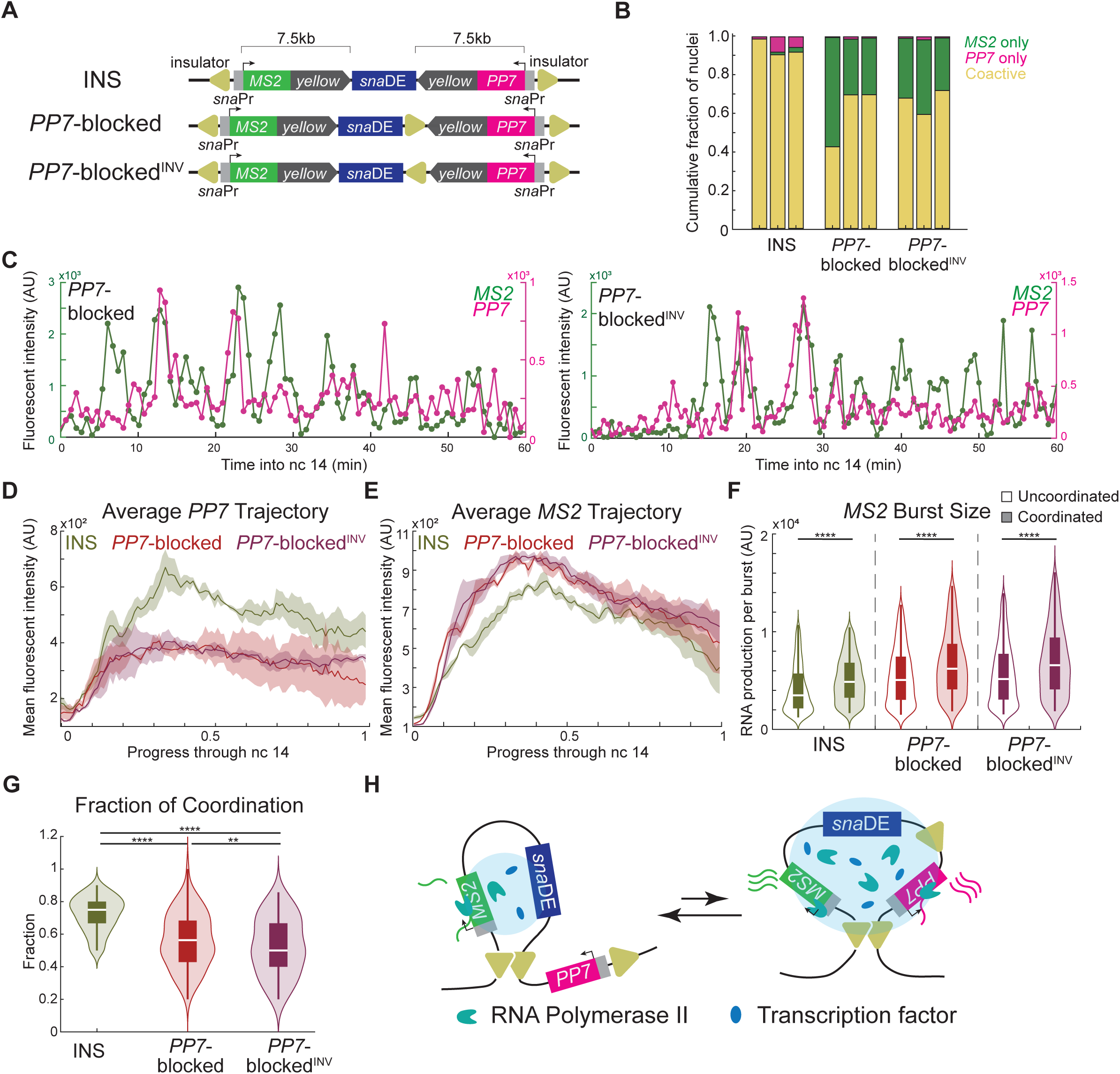
Insulator-mediated looping is dynamic and dominated by distance. (A) Schematic of transgenic constructs containing flanking (INS) and intermediate insulators (PP7-blocked and PP7-blocked^INV^). (B) Bar chart displaying cumulative fraction of nuclei that show only MS2 activity (green), only PP7 activity (magenta) and coactivity (yellow) (N = 3 biological replicates for each construct, one bar per replicate). The presence of an intermediate insulator greatly increases the proportion of MS2-monoactive nuclei, yet coactive nuclei remain a significant fraction. (C) Representative MS2 (green) and PP7 (magenta) trajectories from PP7-Blocked (left) and PP7-Blocked^INV^ (right). (D) Average PP7 trajectory of PP7 active nuclei (N = 3 biological replicates for each construct). The solid line represents the average fluorescence at each time point and the shaded region denotes the standard error. n(INS) = 747, n(PP7-Blocked) = 442, n(PP7-Blocked^INV^) = 684. (E) Average MS2 trajectory of MS2 active nuclei (N = 3 biological replicates for each construct). The solid line represents the average fluorescence at each time point and the shaded region denotes the standard error. n(INS) = 747, n(PP7-Blocked) = 724, n(PP7-Blocked^INV^) = 1001. (F) Violin plot of MS2 burst size for each construct sorted into uncoordinated (unfilled, left) and coordinated (filled, right) bursts. Burst size data was compiled and sorted from three biological replicates. n(INS,Uncoordinated) = 2723, n(INS,Coordinated) = 4384, n(PP7-Blocked,Uncoordinated) = 4876, n(PP7-Blocked,Coordinated) = 1370, n(PP7-Blocked^INV^,Uncoordinated) = 7582, n(PP7-Blocked^INV^,Coordinated) = 1921. (G) Violin plot of Fraction of Coordination. n(INS) = 566, n(PP7-Blocked) = 352, n(PP7-Blocked^INV^) = 541. (H) Illustration of model, showing dynamic looping of system, preferential short-range pairing and promoter synergy during coordinated bursting. **p<0.01, ****p<0.0001 by Wilcoxon rank-sum test (F) or by one-way ANOVA and Tukey-Kramer test (G).

This finding raised a critical question regarding looping dynamics: were these nuclei stably locked into either a coactive or PP7-exclusionary looping state for the entirety of nc 14, or could they oscillate between the two? If looping were static, we would then expect coactive nuclei to assume the system-wide loop and MS2-only monoactive active nuclei to assume a PP7-exclusionary loop for all of nc 14. As such, PP7-active nuclei from both INS and PP7-Blocked should have comparable PP7 transcriptional activities. Conversely, if looping were dynamic, we would expect the alternations in topology to result in distinct PP7 transcriptional behaviors between INS and PP7-Blocked. Analysis of PP7 transcriptional trajectories and averaged fluorescent intensities of PP7-active nuclei revealed a significantly reduced level of PP7 activity in PP7-Blocked compared to INS (Figures 4C, 4D and S4A). Hence, those nuclei that overcame or escaped blocking did so transiently, indicative of dynamic looping, and the changing topology ultimately impaired the interaction of the enhancer with the PP7 promoter. This reduction was largely due to a reduced PP7 bursting frequency (Figure S4B), consistent with the theory that insulators modulate E-P interaction frequency^11^. Critically, the size of the PP7 bursts that did occur was indistinguishable between PP7-Blocked and INS (Figure S4C), indicating that the E-P interaction was just as productive once established. A statistically-similar, burst frequency-driven reduction in PP7 activity was observed in PP7-Blocked^INV^, suggesting that regardless if the intermediate insulator is preferentially pairing with the MS2-or PP7-promoter proximal insulator, the ability of the PP7 promoter to interact with the enhancer was equally hindered (Figure S4B). Similarly, PP7 burst size of PP7-Blocked^INV^ was indistinct from INS and PP7-Blocked (Figure S4C). Therefore, the intermediate insulator, irrespective of relative orientation, dilutes the time the enhancer spends engaged with the PP7 promoter without diminishing PP7’s intrinsic transcriptional capacity.

Intriguingly, we found further insight into dynamic looping from analyzing the transcriptional activity from the unblocked MS2 reporter, whose expression was enhanced upon insertion of an intermediate insulator (Figures 4E and S4D). In contrast to the frequency-driven decrease in PP7 activity, this augmentation was not the product of bursting frequency (Figure S4E). Rather, the MS2 burst size was substantially increased upon blocking the PP7 reporter (Figures 4F and S4F). This unexpected result raises two possibilities: either MS2 bursts became larger because the enhancer remained engaged with the MS2 promoter for longer (a duration-driven effect), or because burst initiation itself became more efficient once PP7 was sequestered, perhaps through redirection of freed transcriptional machinery to the MS2 promoter (an amplitude-driven,’resource opportunism’ effect). These two mechanisms make opposing predictions: opportunism, acting through increased initiation efficiency, should elevate both burst frequency and amplitude; prolonged engagement, by contrast, should increase burst duration without altering amplitude, while leaving frequency unchanged or reduced.

We found that the increased MS2 burst size observed in PP7-blocked constructs was accompanied by a slight but significant reduction in MS2 burst frequency and an increase in MS2 burst duration (Figures S4E-S4G), favoring prolonged enhancer–MS2-promoter engagement as the dominant mechanism. Importantly, this duration-driven interpretation does not require us to specify with which insulator the intermediate pairs. In principle, the intermediate insulator can form loops of differing lengths, including the shorter PP7-isolating loop in which the intermediate pairs with the PP7-flanking insulator. PP7-Blocked behavior thus represents an ensemble average over these competing configurations rather than a single fixed topology. Consistent with the transient escape observed in coactive nuclei (Figure 4C), the MS2 monoactive nuclei from the PP7-Blocked construct displayed a unimodal MS2 output distribution (Figure S4H). If looping were static, we would then expect MS2 monoactive nuclei to resolve into multiple, distinct transcriptional populations, one for each loop geometry. Instead, this unimodal distribution supports a model where individual nuclei rapidly interconvert among competing topologies rather than partitioning into discrete structural states. The two-state modeling presented in the following section independently corroborates this interpretation through its effect on the inferred OFF-state duration.

Notably, when the system-wide loop did form in PP7-Blocked, bringing all three elements into proximity, the consequent coordinated bursting retained its superior size compared to uncoordinated bursting (Figure 4F). These coordinated events likely arise from the same system-wide loop configuration in both PP7-Blocked and PP7-Blocked^INV^, independent of intermediate insulator orientation. This confirms that the fundamental properties of coordinated bursting are preserved regardless of the topological complexity of the system. Quantitatively, among coactive nuclei in PP7-Blocked, both the Fraction of Coordination and the Spearman’s correlation between MS2 and PP7 trajectories were similar to or higher than the uninsulated Dual construct (Figures 4G and S4I), indicating that the underlying synchronizing architecture persists even when the intermediate insulator restricts productive PP7 engagement. Relative to INS, however, the Fraction of Coordination, which is calculated using the less bursty reporter, appeared modestly reduced. We attribute this to a measurement artifact rather than a genuine loss of synchrony. Because MS2 activity was highly elevated, the signal sometimes failed to fall below our bursting threshold between bursting events, thereby obscuring the detection of coordination. The two-state modeling in the following section instead infers promoter activity through a probabilistic framework rather than a single fixed intensity cutoff, providing a more robust measure of coordination in the PP7-Blocked system. Additionally, we observed nearly identical transcriptional behaviors regardless of whether the intermediate insulator was in the forward or reverse orientation (Figures 4D-G and S4A-C). Together, these data demonstrate that insulator-mediated chromatin looping is highly dynamic. The system adopts multiple topological configurations within a single developmental window, each with distinct consequences for transcriptional output (Figure 4H).

### Two-state model of transcription reveals insulators as modulators of burst frequency

Our live imaging data revealed clear differences in transcriptional coordination and output across constructs. However, this coordination is not deterministic. Transcription occurs in stochastic periods of activity and inactivity, a phenomenon known as transcriptional bursting.^35^ This stochasticity complicates efforts to directly infer the underlying promoter kinetics from raw trajectories alone. To infer the rates defining the promoter state transitions, we approximate the transcriptional state to a two-state model in which the gene switches between discrete’ON’ (transcriptionally active) and’OFF’ (transcriptionally inactive) states at mathematically inferred rates (Figures 5A and 5B) (see Methods).^37^ Because the transcriptional state kinetics vary depending on genomic context and nearby regulatory elements, we employed this model to examine how modulating local chromatin organization alters transcriptional bursting dynamics.

**Figure 5.**
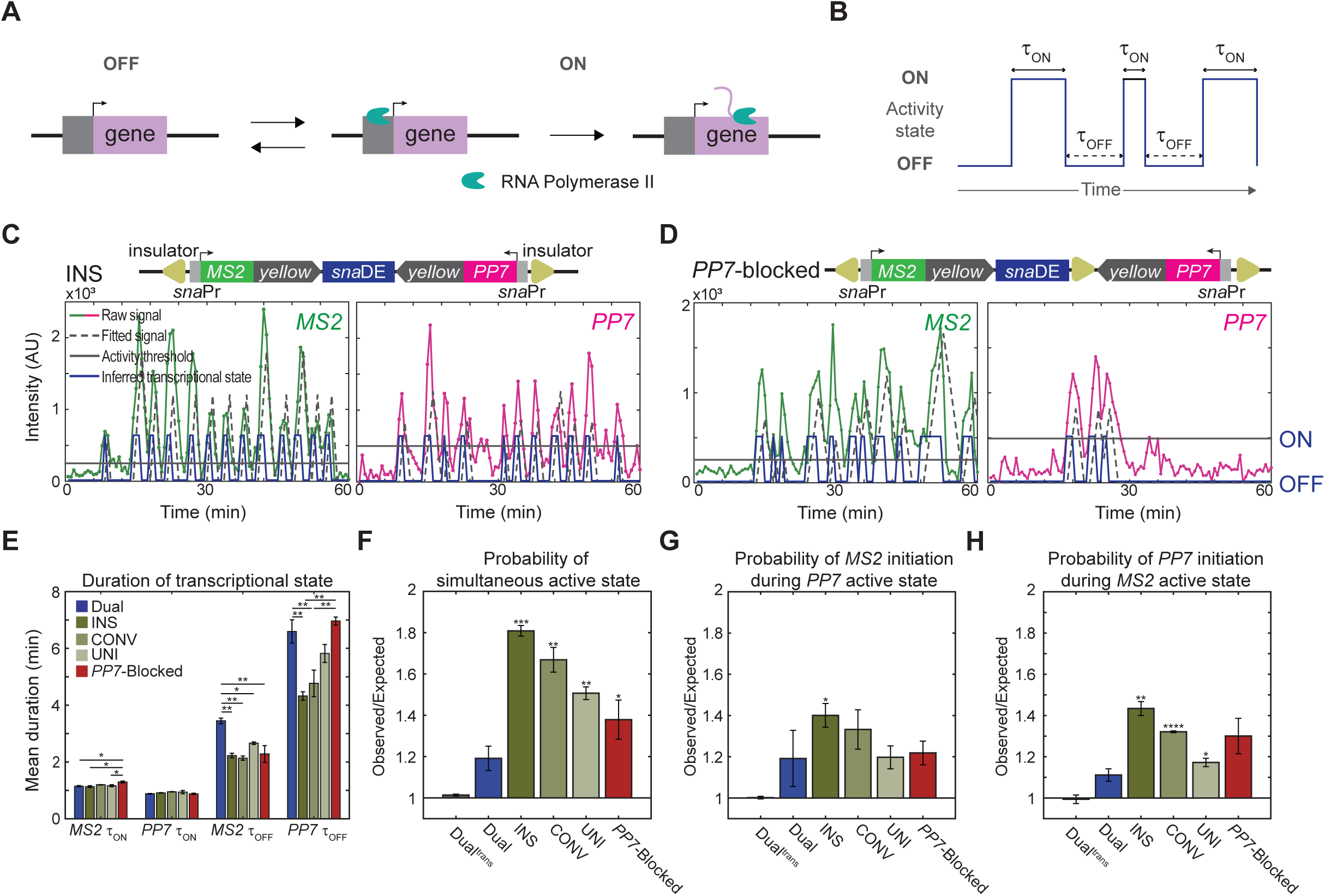
Two-state promoter model reveals insulators as modulators of burst frequency. (A) Schematic of two-state promoter model. (B) Representative trajectory of inferred promoter state, where *τ*_ON_ and *τ*_OFF_ denote periods of activity and inactivity, respectively. (C,D) Raw MS2 and PP7 trajectories (green and magenta lines, respectively), activity threshold (gray line), inferred activity state (blue lines), and fitted signal (dashed lines) plotted across nc 14 for (C) INS and (D) PP7-Blocked constructs. (F) Mean duration of *τ*_ON_ and *τ*_OFF_ for MS2 and PP7 of each construct. (G) Probability of simultaneous MS2 and PP7 active state given either promoter is in active state. (H) Probability of MS2 promoter transitioning from OFF to ON state within 27s time window given PP7 is in active state. (I) Probability of PP7 switching from OFF to ON state within 27s time window given MS2 is in active state. Probabilities are determined per biological replicate from minute 9 to 27 of nc 14. Data are represented as mean ± SEM. *p<0.05, **p<0.01, ***p<0.001, ****p<0.0001 by one-way ANOVA and Tukey-Kramer test (E) or by log-transformed, one-sample t-test (F,G,H).

We applied the two-state model to the uninsulated Dual construct, all flanking insulator configurations (INS, CONV, UNI), and the intermediate insulator configuration (PP7-Blocked). As a negative control, we also modeled the decoupled Dual^trans^ construct, where the reporters no longer share a genomic context or enhancer. The model successfully captured the highly stochastic bursting behavior from all constructs, recapitulating even the rare PP7 activity from the PP7-Blocked construct (Figures 5C and 5D). From the modeled promoter states of individual nuclei, we quantified the duration of active bursting events, *τ*_ON_, and the duration between successive bursts, *τ*_OFF_, for both MS2 and PP7. Overall, their mean values, ⟨*τ*_ON_⟩, ⟨*τ*_OFF_⟩, and their associated variabilities were highly consistent with previous work using sna*DE* (Figures 5E and S5A).^37^ Interestingly, MS2 and PP7 displayed remarkably invariant ⟨*τ*_ON_⟩ durations across the flanking insulator configurations (INS, CONV, UNI) relative to the uninsulated Dual (Figure 5E, ON). This invariance suggests that the increased burst size observed with flanking insulators are not driven by prolonged promoter activation, but rather reflect the enriched transcriptional microenvironment of multi-way E-P co-occupancy. Notably, this pattern differed for the PP7-Blocked construct, in which MS2 ⟨*τ*_ON_⟩ was slightly increased relative to Dual and other flanking insulator configurations (INS, CONV, UNI), consistent with our earlier finding that MS2 burst size augmentation in this construct is driven by prolonged enhancer-MS2 engagement rather than a change in burst frequency (Figure 5E). Similarly, quantification of total activity time (time between the first burst to the last) revealed no significant difference in the presence and absence of insulators, indicating that these elements do not modulate transcriptional onset time (Figure S5B).

In contrast, the mean interburst timing, ⟨*τ*_OFF_⟩, changed significantly across constructs. All insulator-flanked constructs (INS, CONV, and UNI) exhibited a reduced ⟨*τ*_OFF_⟩ relative to the uninsulated Dual construct (Figure 5E, OFF). This finding indicates that insulator-mediated looping lessened the ‘OFF’ time between bursts, i.e. increased the burst frequency. When an additional insulator was positioned between enhancer and PP7 promoter, as is the case with PP7-Blocked, ⟨*τ*_OFF_⟩ increased selectively for PP7 but remained comparable to INS and CONV for MS2, indicating that the intermediate insulator disrupts PP7-enhancer engagement without compromising MS2 burst frequency. Together, this selective lengthening of PP7’s ⟨*τ*_OFF_⟩ without a corresponding cost to MS2 is consistent with dynamic switching between competing loop configurations within a single nuclear cycle (Figure 5E, OFF). Overall, these data provide robust evidence that chromatin insulators primarily function as modulators of burst frequency rather than burst size, aligning with previous findings that topology regulates the probability of E-P interaction.^38^

Next, we leveraged the model to determine whether the promoter states of MS2 and PP7 were temporally correlated. To quantify promoter state coordination, we computed an observed-over-expected (O/E) ratio: the observed probability of a coordinated event divided by the probability expected if the two promoters acted independently (see Methods). By design, an O/E of 1 indicates that the event occurs at random, whereas deviation from 1 signifies non-random coordination. We first tested the simplest prediction of coordination: that the promoters would occupy the ‘ON’ state simultaneously more often than expected by chance (Figures 5F and S5C). However, this test alone cannot fully distinguish genuine coordination from a trivial confound: because both reporters are driven by a strong, highly active *sna*DE enhancer, both promoters spend a large fraction of nc 14 in the’ON’ state regardless of any true coordination. This baseline activity could inflate the apparent frequency of simultaneous’ON’ states even under independent regulation. To control for this, we also assessed whether each promoter showed an increased probability of initiation, i.e. transitioning from’OFF’ to’ON’, specifically while the other promoter was already active (Figures 5G and 5H), a test that isolates genuine coordinated initiation from the shared baseline of high transcriptional activity.

All three O/E metrics yielded consistent trends (Figures 5F-5H and S5C), indicating that the elevated ‘ON’-state overlap observed in Figure 5F reflects genuine coordinated initiation rather than an artifact of shared baseline activity. As expected, the Dual^trans^ control showed baseline, near-random coordination between promoter states. The uninsulated Dual construct showed moderate coordination, whereas all insulated constructs displayed heightened synchronization of transcriptional states. In strong agreement with experimental results, the bidirectional insulator configurations (INS, CONV) yielded greater coordination and correlation than the unidirectional (UNI) configuration. Notably, the intermediate insulator construct, PP7-Blocked, maintained a higher coordination and correlation of promoter states compared to the uninsulated Dual construct. Therefore, while the intermediate insulator may disrupt E-P interaction and productive PP7 transcription, the underlying architecture remains intact and continues to synchronize promoter activity.

Importantly, this coordination is confined to initiation events while the other promoter is ‘ON’. Analysis of alternative transitions, wherein a promoter switches ‘OFF’ while the other is ‘OFF’, switches ‘ON’ when the other is ‘OFF’, or switches ‘OFF’ while the other is ‘ON’, revealed that these events occurred at random or less frequently than expected (Figures S5D-S5F). These results indicate that the coordination of gene activity occurs at transcriptional initiation. Furthermore, this coordination is potentiated in the presence of insulators, suggesting that chromatin architecture fundamentally shapes the probability of E-P interaction. In summary, this modeling approach not only corroborates our empirical findings, but reveals that insulator-mediated synchronization operates specifically at the level of transcriptional initiation, a mechanistic insight unattainable from bulk coactivity measurements alone that further supports insulators as orientation-dependent regulators of E-P interaction frequency.

## Discussion

The coordinated transcriptional activity of neighboring genes is an established and evolutionarily conserved feature of genomic arrangement. While prokaryotes achieve this coordination via polycistronic operons, eukaryotic genomes are believed to direct coexpression through ‘topological operons’ that rely on chromatin architectural elements.^13^ At the molecular level, this coexpression is thought to depend on multi-way E-P interactions, where a single enhancer engages multiple gene targets. For example, the co-initiation of distal paralogues^10^ has been previously documented as one such interaction; however, these findings could arise from stochastic fluctuations or global *trans*-activation events. The exact mechanism governing multi-way E-P interaction remained a central question.

A major contribution of this work is the highly-controlled in *vivo* evidence of how insulator elements modulate real-time transcriptional kinetics. Rather than serving as passive genomic barriers, insulators are active, tunable regulators of local chromatin interactions. We reveal a dual effect of insulator-mediated looping on multi-way E-P interactions. First, the presence of flanking insulators encourages E-P interactions, presumably through a reduced search space, and is consistent with the heightened intra-TAD contact frequency observed in chromatin conformation capture assays.^34^ Second, the insulator-mediated looping biases the system toward simultaneous, rather than sequential, multi-way E-P engagement.

This simultaneity is distinct from other models of E-P communication based on rapid, sequential contacts (e.g., “kiss and kick”). Instead, our real-time trajectories and promoter-state modeling indicate a genuine spatial co-occupancy of enhancer and promoters, in which a shared microenvironment of TFs and transcriptional machinery allows the enhancer to initiate transcription from both promoters at once. This biophysical picture is supported by evidence that coordinated bursting is preceded by greater clustering of transcriptional resources than single-promoter engagements.^36^ Given that burst size correlates with Pol II recruitment and pause-release efficiency,^35^ this local pooling of resources explains why coordinated bursts are larger than uncoordinated ones.

Critically, insulator-mediated looping appears to actively promote this co-occupancy: by increasing the frequency with which the enhancer engages both promoters simultaneously, insulators enrich the local microenvironment with transcriptional resources beyond what either promoter would recruit independently, resulting in larger, more productive bursts from both genes. In contrast to promoter competition, this facilitated proximity and pooling of resources appears highly synergistic in nature such that both promoters are significantly more productive as a consequence of the established local chromatin organization. This active, transcription-dependent view of insulator function echoes recent findings in early *Drosophila* embryos, where an active promoter was found to contribute more to boundary insulation than the insulator-bound region itself, suggesting a functional cross-talk between transcriptional activity and insulator-mediated genome organization.^39^

Importantly, this facilitation is a function of relative insulator orientation. The bidirectional insulator configurations displayed transcriptional behaviors consistent with more stable looping compared to the unidirectional configuration. This disparity is explained by the topological rule of homotypic insulators preferentially pairing in a ‘head-to-head’ manner. When interacting in cis, the oppositely-oriented insulators can more readily form a simple ‘stem loop’, whereas unidirectional insulators require that the chromatin loop back on itself into an entropically-unfavorable ‘circle loop’.^33,40^ Our work provides compelling evidence that this difference in pairing capability translates directly into divergent transcriptional regimes.

Yet even the more stable, bidirectional configurations did not lock the system into a single, fixed topology. By enabling the formation of multiple domains, we highlight that chromatin loops are not static, rigid structures. For instance, monoactive MS2 output in PP7-Blocked formed a single, unimodal distribution rather than the two distinct populations expected from stable, separable loop geometries, indicating that individual nuclei sample multiple configurations within a single nuclear cycle. The highly-dynamic nature of insulator-mediated chromatin looping has real-time effects on transcriptional activity. This is consistent with recent genome-wide quantification of chromatin looping, which found that most loops (including E-P loops) form with low probability and therefore exist only transiently in any given cell, generalizing our live-imaging observations at an individual locus to the genome as a whole.^41^ While similar results of transient insulator behavior have been reported in mammalian systems, such as CTCF-mediated loops in mouse embryonic stem cells,^42^ those studies have not directly linked such dynamics to real-time transcriptional consequences. Our study couples dynamic topology to instantaneous transcriptional activity within living *Drosophila* embryos.

While these findings clarify the relationship between chromatin organization and transcription, several questions remain. First, because we did not directly visualize chromatin conformation, the described topologies have been inferred indirectly from single-nucleus transcriptional trajectories and quantitative two-state modeling of promoter kinetics. While this modeling framework provides a rigorous, quantitative basis for inferring the underlying looping dynamics, direct visualization through simultaneous live imaging of DNA and nascent transcription will be necessary to unambiguously confirm the proposed topological configurations. Second, our functional characterization focused solely on a Su(Hw)-dependent insulator element. Su(Hw) recruits the same architectural proteins (such as Centrosomal protein 190 and an isoform of Mod(mdg4)) as other insulators within the fly, suggesting a conserved mechanism of action.^43,44^ However, confirming if these kinetic rules generalize to other insulators remains an open avenue of inquiry, particularly given evidence that distinct Drosophila insulator proteins contribute to boundary function in a locus-specific, non-redundant manner.^39^ Third, our experiments were restricted to a specific genomic locus during a defined developmental window to minimize confounding variables. We recognize that it limits the generalizability of our findings as they pertain to other genomic loci, cell types and life stages.

Ultimately, this study elucidates another lever of gene regulation in which the 3D architecture of the genome does not merely constrain expression, but rather actively optimizes it. Our work demonstrates that spatial co-occupancy within an enhancer-driven microenvironment allows target promoters to relinquish the limitations of binary competition, achieving a cooperative, orientation-tunable state of synchronous bursting whose frequency is directly sculpted by chromatin topology. As research continues to uncover additional intricacies of higher-order chromatin structure, the principles established here offer a foundational framework for understanding how eukaryotic cells harness dynamic topology to drive robust gene expression throughout development.

## Acknowledgements

We thank Lim lab members and alumni for helpful discussions and feedback on the manuscript. We also thank Paulina Bargallo Gonzalez Lugo for help in cloning and imaging. **Funding:** This work was supported by National Institutes of Health grant R35GM133425 (B.L.) NIH-NIGMS grant R35GM148351(A.S.), and the University of Pennsylvania Fontaine Society (N.B.).

## Declaration of Interests

The authors declare no competing interests.

## Supplemental Movie 1. Live-imaging of Insulated (INS) construct during nc 14

Max-projected imaging of MS2-yellow reporter (left), PP7-yellow reporter (center), and merged (right) from INS strain. Nascent transcripts are labeled with MCP-GFP and PCP-mCherry shown in green and magenta, respectively. His2Av-eBFP2, in blue, is used to visualize nuclei. Images are oriented with anterior to the left. Scale bar represents 10um.

## Supplemental Movie 2. Live-imaging of PP7-Blocked construct during nc 14

Max-projected imaging of MS2-yellow reporter (left), PP7-yellow reporter (center), and merged (right) from PP7-Blocked strain. Nascent transcripts are labeled with MCP-GFP and PCP-mCherry shown in green and magenta, respectively. His2Av-eBFP2, in blue, is used to visualize nuclei. Images are oriented with anterior to the left. Scale bar represents 10um.

## Materials & Methods

### Fly strains generated via PhiC31 Integration

For all experiments, *Drosophila melanogaster* embryos were observed for the entirety of nuclear cycle 14. All plasmids containing our reporters were integrated to the vk33 landing site on the third chromosome (Bloomington Drosophila Stock Center, cat# 9750) via micro-injection performed at BestGene, Inc.^45^ Virgin progeny from the nos>MCP-GFP, PCP-mCherry, His-BFP^46^ and yw cross were crossed with males carrying the MS2 and PP7 reporter genes. Embryos from the cross were imaged under a confocal microscope at room temperature.

### Plasmid construction

All plasmids were generated from the previously published *pbphi-snaPr-MS2-yellow-snaSE-yellow-PP7-snaPr* plasmid.^46^ Its cloning is documented as the following:

pbphi-snaPr-MS2-yellow-sna Shadow-yellow-PP7-snaPr

Plasmid DNAs containing linked MS2-yellow and PP7-yellow reporter genes were created as follows. Towards this end, a DNA fragment containing snaPr-PP7-yellow was purified from pbphi-snaPr-PP7-yellow by digesting with XbaI. The resulting fragment was inserted into the unique XbaI site in pbphi-snaPr-MS2-yellow. Subsequently, a DNA fragment containing the sna shadow enhancer was amplified from genomic DNA using primers (5 ‘-GGG GGA AGC TTG CAT TGA GGT GTTTTG TTG G-3’) and (5 ‘-CCC CCG CTA GCT AAA TTC CGA TTT TTC TTG TCTGGG-3’) and digested with HindIII and NheI. The resulting fragment was inserted between the HindIII and NheI sites of the plasmid.

pbphi-gypsy-snaPr-MS2-yellow-sna Shadow-yellow-PP7-snaPr-gypsy

Insulator sequence was amplified from plasmid DNA using primers (5’-AATT GCGGGCC GCC TGG CCA CGT AAT-3’) and (5’-AATT GCGGCC GCG TTG TTG GTT GGC-3’) and digested with NotI. Insertion of this sequence to the single NotI site upstream of snaPr>MS2-yellow yielded both the forward and reverse orientation. Amplification of gypsy with primers (5’-GGCC TCTAGA CTG GCC ACG TAAT-3’) and (5’-AATT ACT AGT CTG GCC ACG TAA TAA GTG TGC G-3’), digestion with XbaI, and insertion into the XbaI site upstream of snaPr>PP7-yellow yielded both the forward and reverse orientation. Amplifying one end of the insulator with SpeI ligated and randomly abolished one XbaI cut site. Selection for the desired insertion enabled reintroduction of the snaPr>PP7-yellow reporter via XbaI.

pbphi-gypsy-snaPr-MS2-yellow-sna Shadow-gypsy-yellow-PP7-snaPr-gypsy

Insulator sequence was amplified with primers (5’-AATT GCTAGC CTG GCC ACG TAAT-3’) and (5’-AATT GCT AGC GTT GTT GGT TGGC-3’) and digested with NheI. It was inserted to a single NheI cut site within pbphi-gypsy-snaPr-MS2-yellow-sna shadow-yellow-PP7-snaPr-gypsy that exists between sna shadow and snaPr>PP7-yellow. Doing so yielded both orientations of this intermediate insulator.

pbphi-gypsy-snaPr-MS2-yellow-spacer-yellow-PP7-snaPr-gypsy

To assess whether the flanking insulators themselves induced transcriptional activation, we removed the sna shadow enhancer element via restriction enzyme digest of pbphi-gypsy-snaPr-MS2-yellow-sna shadow-yellow-PP7-snaPr-gypsy with HindIII and NheI. The neutral spacer sequence originates from Leyes Porello et al., 2026^47^and was amplified from plasmid DNA using primers (5’-CAA AAC CAT GCG AAG CTC TCT AGC AAG CTT CAT GGT TAT GTC TCC AGA TT-3’) and (5’-CAA AAC CAT GCG AAG CTC TCT AGA GCT AGC GAA ACT TGT ACG TCG TAG AA-3’). It was then digested with HindIII and NheI and inserted into the enhancer-less backbone plasmid.

### Imaging and analysis

#### Live imaging

Yw; nanos>SV40NLS-mCherry-PCP, His2Av-eBFP2; nanos>MCP-GFP virgins were crossed with males carrying the MS2/PP7 reporter genes. Embryos from the cross were dechorionated and mounted in Halocarbon oil 27 (Sigma) between a semipermeable membrane (Sarstedt) and coverslip (18 mm × 18 mm). Embryos were imaged using a Zeiss LSM800 and Plan-Apochromat 40x/1.3 numerical aperture (N.A.) oil-immersion objective. A single image was 512 x 512 pixels. Images were created by taking a maximum projection from a stack of 14 images separated by 0.75 μm, and the final time resolution was 27s per frame. Image capture started from the beginning of nc 14 mitosis to gastrulation. Images were captured in 16-bit. Laser power was consistent across imaging of all constructs.

#### Image analysis

Segmentation and tracking of nuclei were carried out following the methodology outlined in Keller et al., 2020.^48^ Every *Drosophila* strain used in this research expressed His2Av-eBFP2 to label nuclei. For live image processing and nuclei segmentation, a Gaussian filter was initially applied to the nuclei-labeled channels to smooth images. This was followed by top-hat filtering and contrast-limited adaptive histogram equalization to correct for uneven illumination and to improve image contrast. The processed images were then binarized using a threshold determined by Otsu’s method. Manual corrections were made to the binary images of each frame to ensure segmentation accuracy. Each nucleus identified in the segmentation was assigned an index, and the x and y coordinates of all constituent pixels, as well as the center of mass, were documented. A custom MATLAB script was used to track segmented nuclei from nuclear cycle 14 up to the beginning of gastrulation. To track each indexed nucleus, the Euclidean distance between a nucleus and all nuclei in the following frame was computed and compared. Nucleus pairs with the smallest Euclidean distance, provided it was less than 10 pixels between consecutive frames, were matched and considered to represent the same nucleus over time. If the minimum distance exceeded 10 pixels, the nucleus was classified as having no corresponding lineage, due to partial or complete movement out of the imaging frame. MS2 and PP7 signal intensities were extracted by taking the average of the highest two pixels from each nucleus using a mask created by segmenting the embryo.

#### Quantification

Background subtraction for the MS2 and PP7 channels were performed by subtracting the respective channel’s mean intensity of initial non-transcribing signals within all nuclei of an individual embryo. Then, each MS2 and PP7 trace was individually zeroed by subtracting its minimum value from all points. The mRNA output was measured by taking the area under the curve of each MS2 and PP7 trajectory over the entire nc 14 time course. Burst size was calculated by using a threshold to identify the onset and offset of individual bursting events, linearly interpolating 1000 points between onset and offset and then integrating beneath this interpolated curvature. Burst frequency was determined by counting the number of bursting onset events per nucleus throughout nc 14. Burst amplitude was selected as the maximum value between onset and offset of individual bursting events, i.e. the peak of each burst. Duration was the number of frames between onset and offset, using the 1000 point interpolated curve to determine more precise durations. The Spearman’s Correlation Coefficient was determined from the following equation:

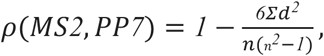

where d is the difference in rank between the same time point in the trace and n is the number of time points in the trace. The Fraction of Coordination was determined by first categorizing individual bursting events into either ‘Uncoordinated’ or ‘Coordinated’. Designation of ‘Coordinated’ bursts required that the onset of bursting for both reporter genes occurred within a two minute window. If not, then it was designated as ‘Uncoordinated’. Then, for each individual coactive nucleus, the number of ‘Coordinated’ bursts was divided by the total number of bursts to calculate the Fraction of Coordination. To keep the metric consistent across all constructs regardless of the relative activity of the reporter genes, the Fraction of Coordination was calculated and reported for whichever reporter gene had a lower burst frequency.

### Mathematical modeling and transcriptional state inference

#### Inference of Transcriptional States

To infer the binary transcriptional states (ON/OFF) from single-cell MS2 and PP7 fluorescence trajectories, we applied a customized transcriptional state inference algorithm based on a deconvolution method.^37^ First, raw fluorescence signals were baseline-corrected by subtracting the minimum signal value for each nucleus and smoothed using a flat-top window convolution (window size = 7 frames or 3 min) to reduce high-frequency noise. The continuous transcription rate was approximated using the discrete temporal derivative of the smoothed signal, normalized by a given loading rate (linear slope of the accumulation signal) and the experimental time step. To exclude fluctuations in the basal level, only signal values above a defined fluorescent threshold were retained. The resulting continuous transcription rate array was subsequently smoothed using a boxcar convolution (window size = 2 frames or 54 sec) and discretized into binary promoter states, where values exceeding an activity threshold were assigned as the active (ON) state, and all others as the inactive (OFF) state.

To estimate the optimal loading rate and transcriptional binary state for each nucleus, we employed a grid-search optimization, evaluating loading rates between a wide range that was chosen such that all nuclei fit in that range. For each tested rate, an idealized theoretical fluorescence trajectory was reconstructed from the inferred binary states. The optimal loading rate for a given nucleus was selected by minimizing the sum of squared residuals between this theoretical trajectory and the smoothed experimental data. Finally, the durations of the resulting ON and OFF intervals were extracted to calculate key kinetic parameters, including ⟨*τ*_ON_⟩, ⟨*τ*_OFF_⟩ and total activity time. The states before the first burst and after the last burst were neglected for the estimation of the burst statistics (mean duration and variability) to account for global transcriptional inactivity at the beginning and end of nc 14.

#### Calculation of Joint Probabilities and Association Ratios

To evaluate the co-occurrence of transcriptional states between the MS2 and PP7 reporters, we calculated empirical joint probabilities and their deviation from random expectation. The analysis was restricted to the steady-state window (minute 9 to 27 of nc 14), in which the average transcriptional state was approximately steady and far from the trivial state (promoters OFF) at the beginning and the end of the nuclear cycle. This analysis was performed for each individual nucleus in each biological replicate before being aggregated. First, the marginal probabilities of each reporter P(MS2_ON_) or P(MS2_OFF_) and P(PP7_ON_) or P(PP7_OFF_) were determined from the relative frequency of each state by calculating the number of nuclei in the given transcriptional state divided by the total number of nuclei in the given embryo. To estimate these statistics, we treated states in different frames for the same nucleus as independent. Next, the empirical joint probabilities for all four possible state combinations P(MS2_ON_,PP7_OFF_), P(MS2_OFF_,PP7_OFF_), P(MS2_OFF_,PP7_ON_) and P(MS2_ON_,PP7_ON_) were derived by calculating the frequency of simultaneous state occurrences within the same time frame. In total, each nucleus contributes 40 points (40 frames spanning minute 9 to 27 of nc 14) to the pooled dataset. To determine if the observed co-occurrences occurred more or less frequently than expected by random chance, an O/E ratio was computed for each combination. This ratio was defined as the empirical joint probability divided by the product of the corresponding marginal probabilities:

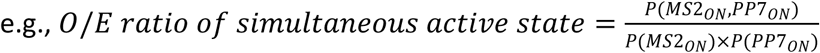

An O/E ratio of 1 indicates independent regulation, while values greater or less than 1 indicate positive or negative association, respectively.

#### Estimation of Conditional Switching Probabilities

To quantify the influence of one promoter’s transcriptional state on the switching dynamics of the other, we calculated the O/E ratio of the conditional switching probability to the marginal (random) switching probability. The temporal analysis was restricted to the steady-state window spanning minute 9 to 27 of nc 14. First, we computed the baseline marginal transition probabilities for each reporter gene (MS2 and PP7) switching between active (ON) and inactive (OFF) states. For instance, the marginal probability of MS2 activating was calculated as the total number of OFF→ON transitions divided by the total number of frames where MS2 was initially OFF. This gives us an estimator of the probability P(MS2_OFF→ON_). We then calculated the conditional switching probabilities, constrained by the concurrent state of the other reporter – for example, the probability of MS2 turning ON only when PP7 is ON, P(MS2_OFF→ON_|PP7_ON_), as the number of MS2 OFF→ON transitions occurring while PP7 was ON, divided by the number of frames in which MS2 was OFF and PP7 was ON. All the switching events of the studied nuclei were gathered and treated as independent events. To estimate uncertainty and preserve biological variability, these ratios were computed independently for each construct.

#### Estimation of the correlation coefficient of transcriptional states

To quantify the coordination between the MS2 and PP7 transcriptional states, we calculated a discrete correlation metric for each biological replicate. First, the inferred transcriptional trajectories were restricted to a steady-state window (min 9 to 27 of nc 14) and treated as explicit binary variables, where a state of 1 represented active transcription (ON) and 0 represented an inactive promoter (OFF). To generate correlation estimates and 95% confidence intervals, we employed a statistical bootstrapping approach with 500 resampling iterations per replicate. For each iteration, the temporal data were randomly resampled with replacement, and the empirical probabilities of MS2 being active (P(MS2_ON_)), PP7 being active (P(PP7_ON_)), and their simultaneous co-occurrence (P(MS2_ON_,PP7_ON_)) were computed as the arithmetic means of the resampled binary arrays. The correlation coefficient (R) for each bootstrap sample was calculated as the covariance of the two discrete states divided by the product of their standard deviations:

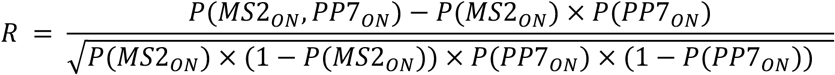

After the 500 bootstrapping iterations, the median of the resulting bootstrap distribution (the 50th percentile) was taken as the central correlation estimate for that specific replicate. The 2.5th and 97.5th percentiles were extracted as the lower and upper bounds of the 95% confidence intervals.

## Notes

### Competing Interest Statement

The authors have declared no competing interest.

## References

1. Levine, M. (2010). Transcriptional Enhancers in Animal Development and Evolution. Curr. Biol. 20, R754–R763. 10.1016/j.cub.2010.06.070.

2. Fudenberg, G., Imakaev, M., Lu, C., Goloborodko, A., Abdennur, N., and Mirny, L.A. (2016). Formation of Chromosomal Domains by Loop Extrusion. Cell Rep. 15, 2038–2049. 10.1016/j.celrep.2016.04.085.

3. Du, M., Stitzinger, S.H., Spille, J.-H., Cho, W.-K., Lee, C., Hijaz, M., Quintana, A., and Cissé, I.I. (2024). Direct observation of a condensate effect on super-enhancer controlled gene bursting. Cell 187, 331–344.e17. 10.1016/j.cell.2023.12.005.

4. Chen, H., Levo, M., Barinov, L., Fujioka, M., Jaynes, J.B., and Gregor, T. (2018). Dynamic interplay between enhancer-promoter topology and gene activity. Nat. Genet. 50, 1296–1303. 10.1038/s41588-018-0175-z.

5. Deng, H., Valenti, P., Payre, F., and Lim, B. (2025). 3D dynamics of *trans* enhancer-promoter interactions in living *Drosophila embryos* reveals spatiotemporal thresholds for transcription activation. Preprint at Developmental Biology, 10.1101/2025.03.17.643800.

6. Benabdallah, N.S., Williamson, I., Illingworth, R.S., Kane, L., Boyle, S., Sengupta, D., Grimes, G.R., Therizols, P., and Bickmore, W.A. (2019). Decreased Enhancer-Promoter Proximity Accompanying Enhancer Activation. Mol. Cell 76, 473–484.e7. 10.1016/j.molcel.2019.07.038.

7. Alexander, J.M., Guan, J., Li, B., Maliskova, L., Song, M., Shen, Y., Huang, B., Lomvardas, S., and Weiner, O.D. (2019). Live-cell imaging reveals enhancer-dependent Sox2 transcription in the absence of enhancer proximity. eLife 8, e41769. 10.7554/eLife.41769.

8. Harding, K., Hoey, T., Warrior, R., and Levine, M. (1989). Autoregulatory and gap gene response elements of the even-skipped promoter of Drosophila. EMBO J. 8, 1205–1212. 10.1002/j.1460-2075.1989.tb03493.x.

9. Grosveld, F., van Assendelft, G.B., Greaves, D.R., and Kollias, G. (1987). Position-independent, high-level expression of the human beta-globin gene in transgenic mice. Cell 51, 975–985. 10.1016/0092-8674(87)90584-8.

10. Levo, M., Raimundo, J., Bing, X.Y., Sisco, Z., Batut, P.J., Ryabichko, S., Gregor, T., and Levine, M.S. (2022). Transcriptional coupling of distant regulatory genes in living embryos. Nature 605, 754–760. 10.1038/s41586-022-04680-7.

11. Fukaya, T., Lim, B., and Levine, M. (2016). Enhancer Control of Transcriptional Bursting. Cell 166, 358–368. 10.1016/j.cell.2016.05.025.

12. Sanyal, A., Lajoie, B.R., Jain, G., and Dekker, J. (2012). The long-range interaction landscape of gene promoters. Nature 489, 109–113. 10.1038/nature11279.

13. Batut, P.J., Bing, X.Y., Sisco, Z., Raimundo, J., Levo, M., and Levine, M.S. (2022). Genome organization controls transcriptional dynamics during development. Science 375, 566–570. 10.1126/science.abi7178.

14. Majumder, P., and Cai, H.N. (2003). The functional analysis of insulator interactions in the Drosophila embryo. Proc. Natl. Acad. Sci. U. S. A. 100, 5223–5228. 10.1073/pnas.0830190100.

15. Fujioka, M., Mistry, H., Schedl, P., and Jaynes, J.B. (2016). Determinants of Chromosome Architecture: Insulator Pairing in cis and in trans. PLOS Genet. 12, e1005889. 10.1371/journal.pgen.1005889.

16. Bing, X., Ke, W., Fujioka, M., Kurbidaeva, A., Levitt, S., Levine, M., Schedl, P., and Jaynes, J.B. (2024). Chromosome structure in Drosophila is determined by boundary pairing not loop extrusion. eLife 13, RP94070. 10.7554/eLife.94070.

17. Li, Y., Haarhuis, J.H.I., Sedeño Cacciatore, Á., Oldenkamp, R., van Ruiten, M.S., Willems, L., Teunissen, H., Muir, K.W., de Wit, E., Rowland, B.D., et al. (2020). The structural basis for cohesin-CTCF-anchored loops. Nature 578, 472–476. 10.1038/s41586-019-1910-z.

18. Kellum, R., and Schedl, P. (1992). A Group of scs Elements Function as Domain Boundaries in an Enhancer-Blocking Assay. Mol. Cell. Biol. 12, 2424–2431. 10.1128/mcb.12.5.2424-2431.1992.

19. Gaszner, M., Vazquez, J., and Schedl, P. (1999). The Zw5 protein, a component of the scs chromatin domain boundary, is able to block enhancer-promoter interaction. Genes Dev. 13, 2098–2107. 10.1101/gad.13.16.2098.

20. Recillas-Targa, F., Bell, A.C., and Felsenfeld, G. (1999). Positional enhancer-blocking activity of the chicken beta-globin insulator in transiently transfected cells. Proc. Natl. Acad. Sci. U. S. A. 96, 14354–14359. 10.1073/pnas.96.25.14354.

21. Scott, K.C., Taubman, A.D., and Geyer, P.K. (1999). Enhancer blocking by the Drosophila gypsy insulator depends upon insulator anatomy and enhancer strength. Genetics 153, 787– 798. 10.1093/genetics/153.2.787.

22. Perry, M.W., Boettiger, A.N., Bothma, J.P., and Levine, M. (2010). Shadow enhancers foster robustness of Drosophila gastrulation. Curr. Biol. CB 20, 1562–1567. 10.1016/j.cub.2010.07.043.

23. Bertrand, E., Chartrand, P., Schaefer, M., Shenoy, S.M., Singer, R.H., and Long, R.M. (1998). Localization of ASH1 mRNA particles in living yeast. Mol. Cell 2, 437–445. 10.1016/s1097-2765(00)80143-4.

24. Garcia, H.G., Tikhonov, M., Lin, A., and Gregor, T. (2013). Quantitative Imaging of Transcription in Living Drosophila Embryos Links Polymerase Activity to Patterning. Curr. Biol. 23, 2140–2145. 10.1016/j.cub.2013.08.054.

25. Vera, M., Tutucci, E., and Singer, R.H. (2019). Imaging Single mRNA Molecules in Mammalian Cells Using an Optimized MS2-MCP System. Methods Mol. Biol. 2038, 3–20. 10.1007/978-1-4939-9674-2_1.

26. Venken, K.J.T., He, Y., Hoskins, R.A., and Bellen, H.J. (2006). P[acman]: A BAC Transgenic Platform for Targeted Insertion of Large DNA Fragments in *D. melanogaster*. Science 314, 1747–1751. 10.1126/science.1134426.

27. Cubeñas-Potts, C., Rowley, M.J., Lyu, X., Li, G., Lei, E.P., and Corces, V.G. (2017). Different enhancer classes in Drosophila bind distinct architectural proteins and mediate unique chromatin interactions and 3D architecture. Nucleic Acids Res. 45, 1714–1730. 10.1093/nar/gkw1114.

28. Yokoshi, M., Segawa, K., and Fukaya, T. (2020). Visualizing the Role of Boundary Elements in Enhancer-Promoter Communication. Mol. Cell 78, 224–235.e5. 10.1016/j.molcel.2020.02.007.

29. Foe, V.E., and Alberts, B.M. (1983). Studies of nuclear and cytoplasmic behaviour during the five mitotic cycles that precede gastrulation in Drosophila embryogenesis. J. Cell Sci. 61, 31–70. 10.1242/jcs.61.1.31.

30. Holdridge, C., and Dorsett, D. (1991). Repression of hsp70 heat shock gene transcription by the suppressor of hairy-wing protein of Drosophila melanogaster. Mol. Cell. Biol. 11, 1894– 1900. 10.1128/mcb.11.4.1894-1900.1991.

31. West, A.G., Gaszner, M., and Felsenfeld, G. (2002). Insulators: many functions, many mechanisms. Genes Dev. 16, 271–288. 10.1101/gad.954702.

32. Fujioka, M., Ke, W., Schedl, P., and Jaynes, J.B. (2025). The *homie* insulator has sub-elements with different insulating and long-range pairing properties. GENETICS 229, iyaf032. 10.1093/genetics/iyaf032.

33. Kyrchanova, O., Chetverina, D., Maksimenko, O., Kullyev, A., and Georgiev, P. (2008). Orientation-dependent interaction between Drosophila insulators is a property of this class of regulatory elements. Nucleic Acids Res. 36, 7019–7028. 10.1093/nar/gkn781.

34. Rao, S.S.P., Huntley, M.H., Durand, N.C., Stamenova, E.K., Bochkov, I.D., Robinson, J.T., Sanborn, A.L., Machol, I., Omer, A.D., Lander, E.S., et al. (2014). A 3D Map of the Human Genome at Kilobase Resolution Reveals Principles of Chromatin Looping. Cell 159, 1665–1680. 10.1016/j.cell.2014.11.021.

35. Leyes Porello, E.A., Trudeau, R.T., and Lim, B. (2023). Transcriptional bursting: stochasticity in deterministic development. Development 150, dev201546. 10.1242/dev.201546.

36. Kawasaki, K., and Fukaya, T. (2023). Functional coordination between transcription factor clustering and gene activity. Mol. Cell 83, 1605–1622.e9. 10.1016/j.molcel.2023.04.018.

37. Nieto, C., Vahdat, Z., Lim, B., and Singh, A. (2025). Regulation of Transcriptional Bursting and Spatial Patterning in Early Drosophila Embryo Development. Preprint, 10.7554/eLife.107610.1.

38. Doyle, B., Fudenberg, G., Imakaev, M., and Mirny, L.A. (2014). Chromatin Loops as Allosteric Modulators of Enhancer-Promoter Interactions. PLoS Comput. Biol. 10, e1003867. 10.1371/journal.pcbi.1003867.

39. Cavalheiro, G.R., Girardot, C., Viales, R.R., Pollex, T., Cao, T.B.N., Lacour, P., Feng, S., Rabinowitz, A., and Furlong, E.E.M. (2023). CTCF, BEAF-32, and CP190 are not required for the establishment of TADs in early *Drosophila* embryos but have locus-specific roles. Sci. Adv. 9, eade1085. 10.1126/sciadv.ade1085.

40. Ke, W., Fujioka, M., Schedl, P., and Jaynes, J.B. (2024). Stem-loop and circle-loop TADs generated by directional pairing of boundary elements have distinct physical and regulatory properties. eLife 13, RP94114. 10.7554/eLife.94114.3.

41. Jusuf, J.M., Yang, J.H., Toppen, J., Grosse-Holz, S., Gabriele, M., Mach, P., Flyamer, I.M., Zechner, C., Giorgetti, L., Mirny, L.A., et al. (2026). Genome-wide absolute quantification of chromatin looping. Nat. Struct. Mol. Biol. 33, 1105–1114. 10.1038/s41594-026-01819-2.

42. Gabriele, M., Brandão, H.B., Grosse-Holz, S., Jha, A., Dailey, G.M., Cattoglio, C., Hsieh, T.-H.S., Mirny, L., Zechner, C., and Hansen, A.S. (2022). Dynamics of CTCF-and cohesin-mediated chromatin looping revealed by live-cell imaging. Science 376, 496–501. 10.1126/science.abn6583.

43. Gerasimova, T.I., Gdula, D.A., Gerasimov, D.V., Simonova, O., and Corces, V.G. (1995). A drosophila protein that imparts directionality on a chromatin insulator is an enhancer of position-effect variegation. Cell 82, 587–597. 10.1016/0092-8674(95)90031-4.

44. Pai, C.-Y., Lei, E.P., Ghosh, D., and Corces, V.G. (2004). The Centrosomal Protein CP190 Is a Component of the gypsy Chromatin Insulator. Mol. Cell 16, 737–748. 10.1016/j.molcel.2004.11.004.

45. Fish, M.P., Groth, A.C., Calos, M.P., and Nusse, R. (2007). Creating transgenic Drosophila by microinjecting the site-specific φC31 integrase mRNA and a transgene-containing donor plasmid. Nat. Protoc. 2, 2325–2331. 10.1038/nprot.2007.328.

46. Lim, B., Heist, T., Levine, M., and Fukaya, T. (2018). Visualization of Transvection in Living Drosophila Embryos. Mol. Cell 70, 287–296.e6. 10.1016/j.molcel.2018.02.029.

47. Leyes Porello, E.A., Wu, J., Fallacaro, S., Dreispiel Juan, E., Trudeau, R.T., Vidal, J.A., Lin, S., Nieto, C., Singh, A., Mir, M., et al. (2026). Enhancer placement impacts transcriptional dynamics in Drosophila embryos. Nat. Commun. 17, 6155. 10.1038/s41467-026-72618-y.

48. Keller, S.H., Jena, S.G., Yamazaki, Y., and Lim, B. (2020). Regulation of spatiotemporal limits of developmental gene expression via enhancer grammar. Proc. Natl. Acad. Sci. 117, 15096–15103. 10.1073/pnas.1917040117.

