## Supplemental Figures for "Insulators As Dynamic, Tunable Regulators of Enhancer-Promoter Coordination in Living *Drosophila* Embryos"

**Supplemental figure 1**

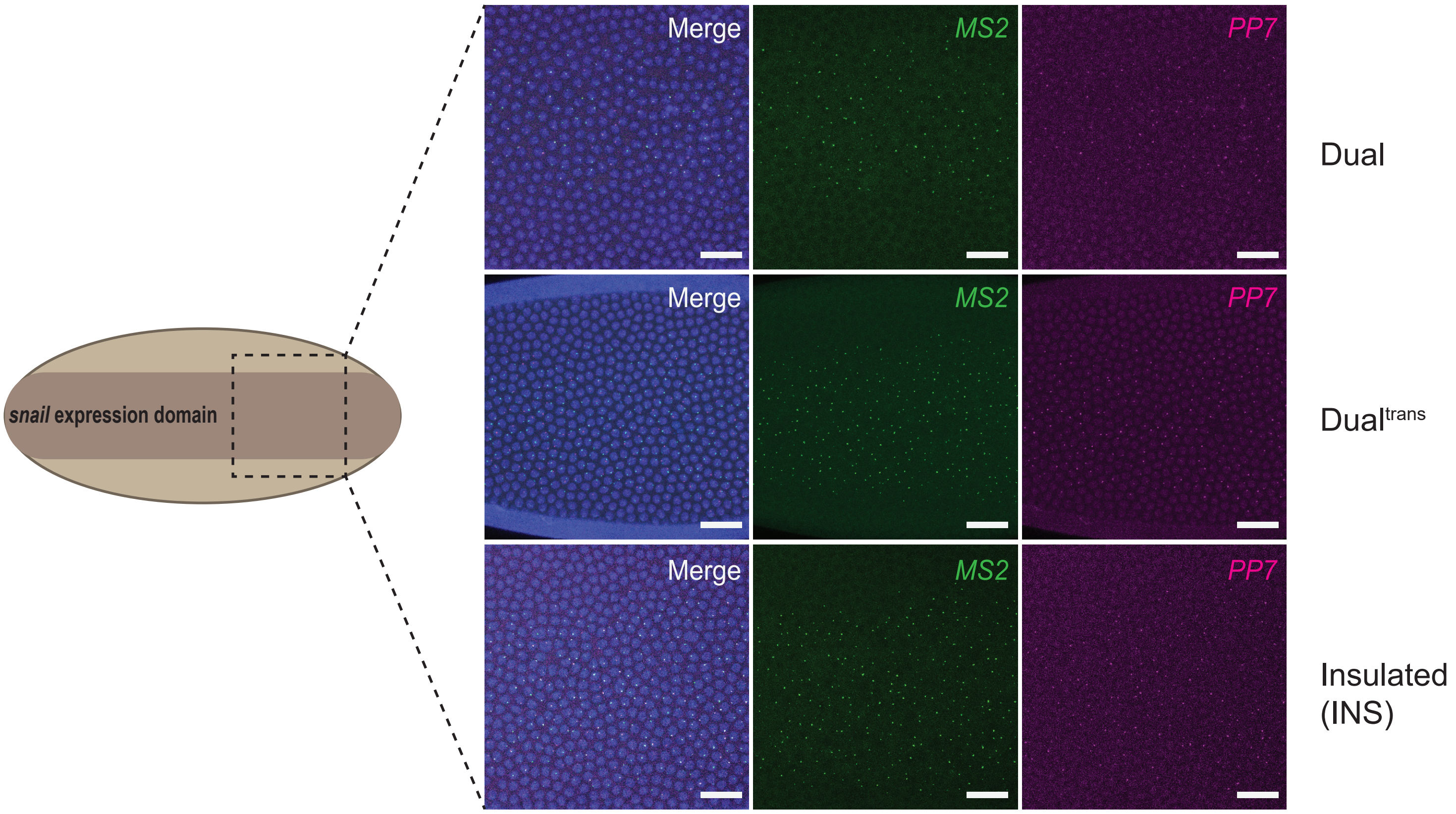


**Supplemental Figure 1.**
(Left) Illustration of *Drosophila* embryo with darkened region representing *snail* expression domain.
(Right) Snapshots from live-imaging of Dual (top), Dual$^{trans}$ (middle) and Insulated (bottom).
Scale bar (white rectangle) represents 20µm.

### Supplemental figure 2

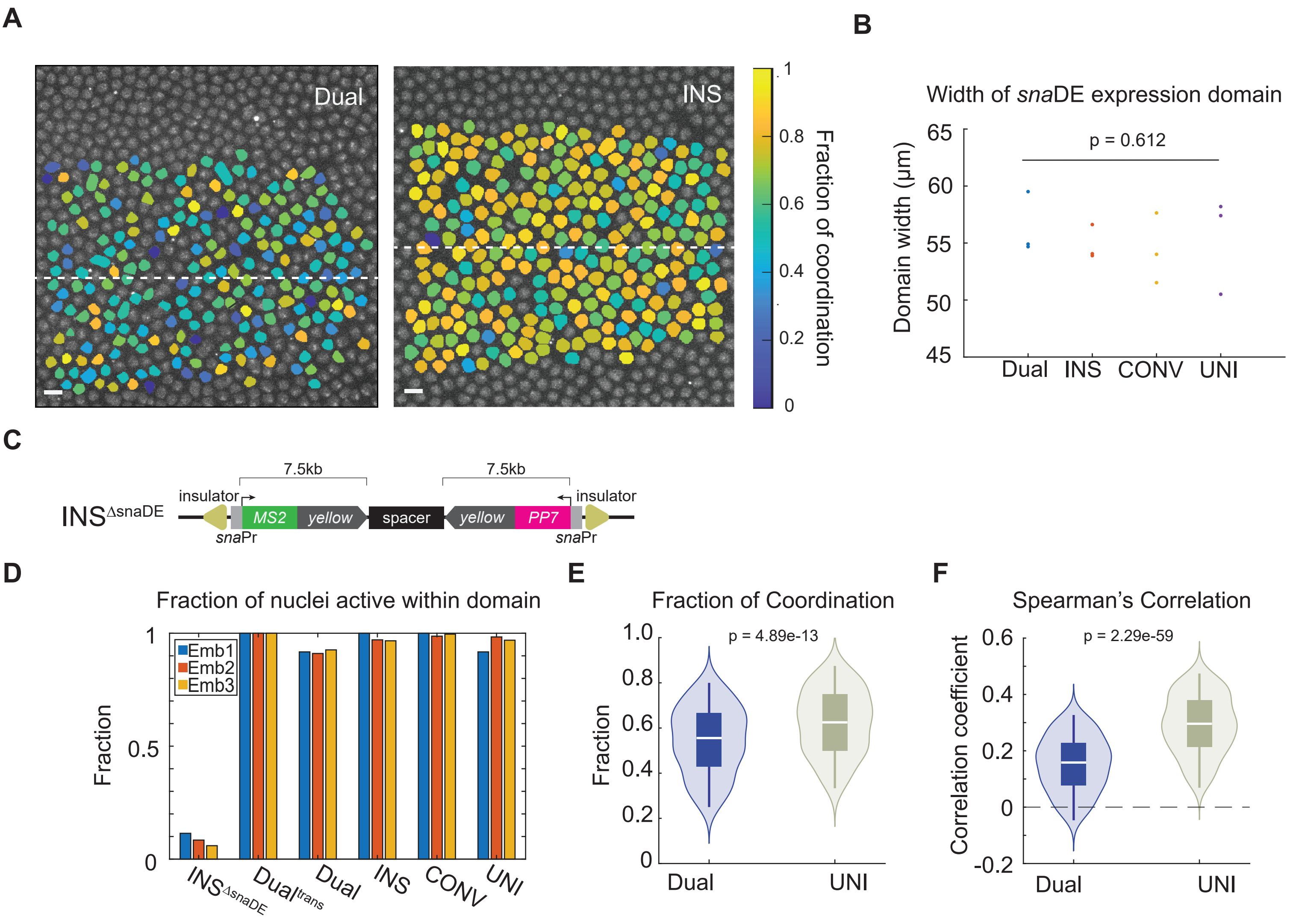


**Supplemental Figure 2.**

(A) False-colored snapshot from imaging depicting Fraction of Coordination of each coactive nucleus in the pictured embryo. White dashed lines represent the ventral midline of the embryo. Scale bar represents 5μm.

(B) Plot of width of expression domain. Each dot represents the average width of the expression domain from four time points within a single biological replicate.

(C) Schematic of enhancerless control construct, $INS^{\Delta snaDE}$.

(D) Bar plot showing the fraction of active nuclei within the domain of expression. Each bar represents a single biological replicate. Replacement of enhancer with neutral spacer sequence yields little to no transcriptional activity.

(E) Violin plot of Fraction of Coordination of Dual and UNI. UNI is significantly higher.

(F) Violin plot of Spearman's Correlation analysis of Dual and UNI, showing heightened correlation in UNI.

Data sets are compiled from three biological replicates. The number of data points is as follows: n(Dual) = 361, n(UNI) = 507.

P-values determined by one-way ANOVA (B) and Wilcoxon rank-sum test (D,E).

### Supplemental figure 3

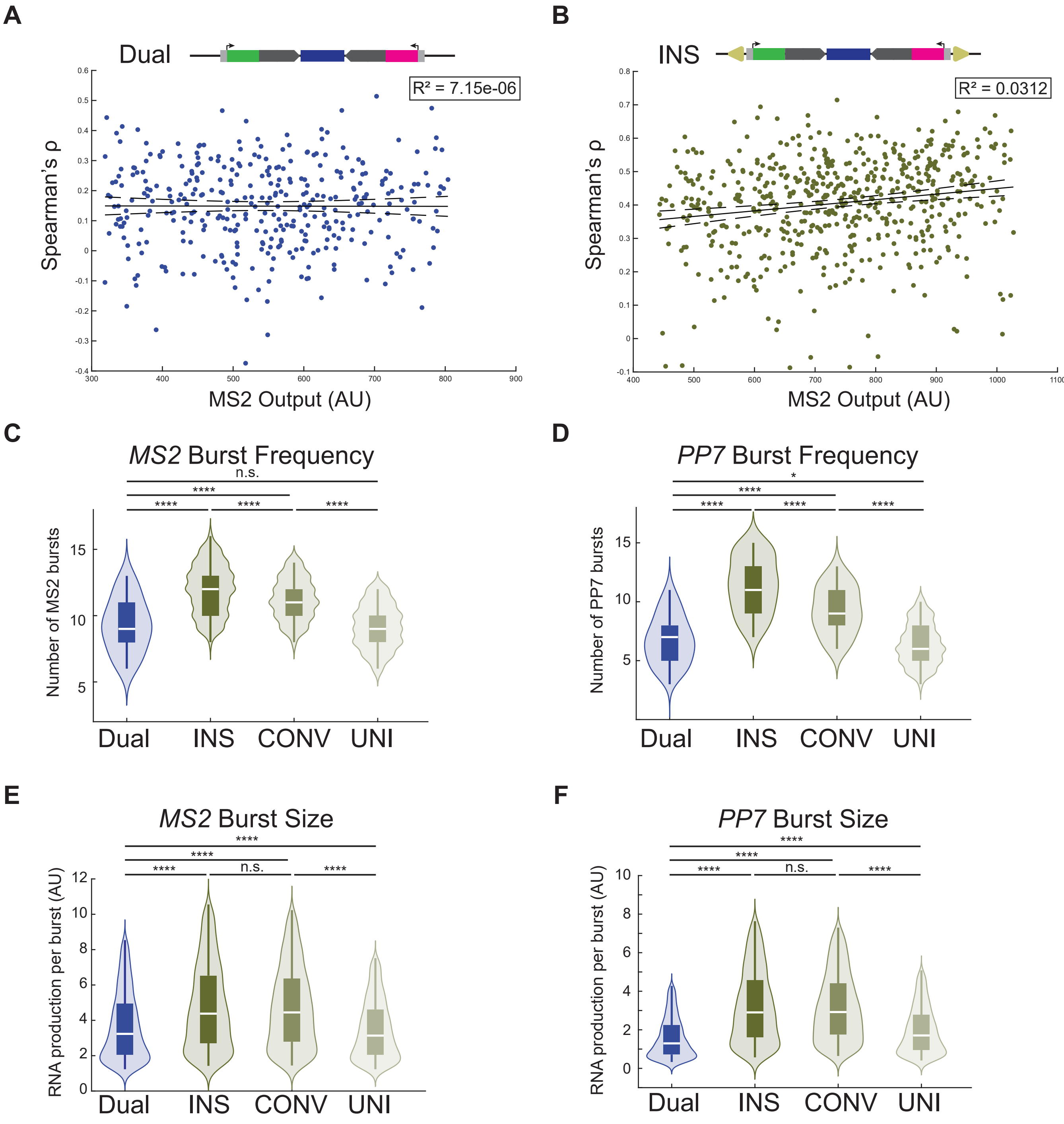


**Supplemental Figure 3.**
(A,B) Scatter plot of *MS2* output versus Spearman's correlation coefficient of individual coactive nuclei from (A) Dual and (B) INS. *MS2* output was calculated from the middle third of nc 14, when most nuclei are active. Linear regression analysis shows little to no correlation. $R^2$ values are displayed. (N = 3 biological replicates, n(Dual) = 361, n(INS) = 566 nuclei).
(C,D) Violin plots of (C) *MS2* and (D) *PP7* burst frequency of Dual, INS, CONV, and UNI. (N = 3 biological replicates, n(Dual,*MS2*) = 457, n(INS,*MS2*) = 573, n(CONV,MS2) = 701, n(UNI,*MS2*) = 587, n(Dual,*PP7*) = 432, n(INS,*PP7*) = 598, n(CONV,*PP7*) = 692, n(UNI,*PP7*) = 617 nuclei).
(E,F) Violin plots of (E) *MS2* and (F) *PP7* burst size of Dual, INS, CONV and UNI. (N = 3 biological replicates, n(Dual,*MS2*) = 4938, n(INS,*MS2*) = 7106, n(CONV,*MS2*) = 7989, n(UNI,*MS2*) = 6454, n(Dual,*PP7*) = 3529, n(INS,*PP7*) = 6889, n(CONV,*PP7*) = 6544, n(UNI,*PP7*) = 4433 bursting events).
n.s. not significant, *p<0.05, **p<0.01, ****p<0.0001 by one-way ANOVA and Tukey-Kramer test (C,D,E,F).

**Supplemental figure 4**

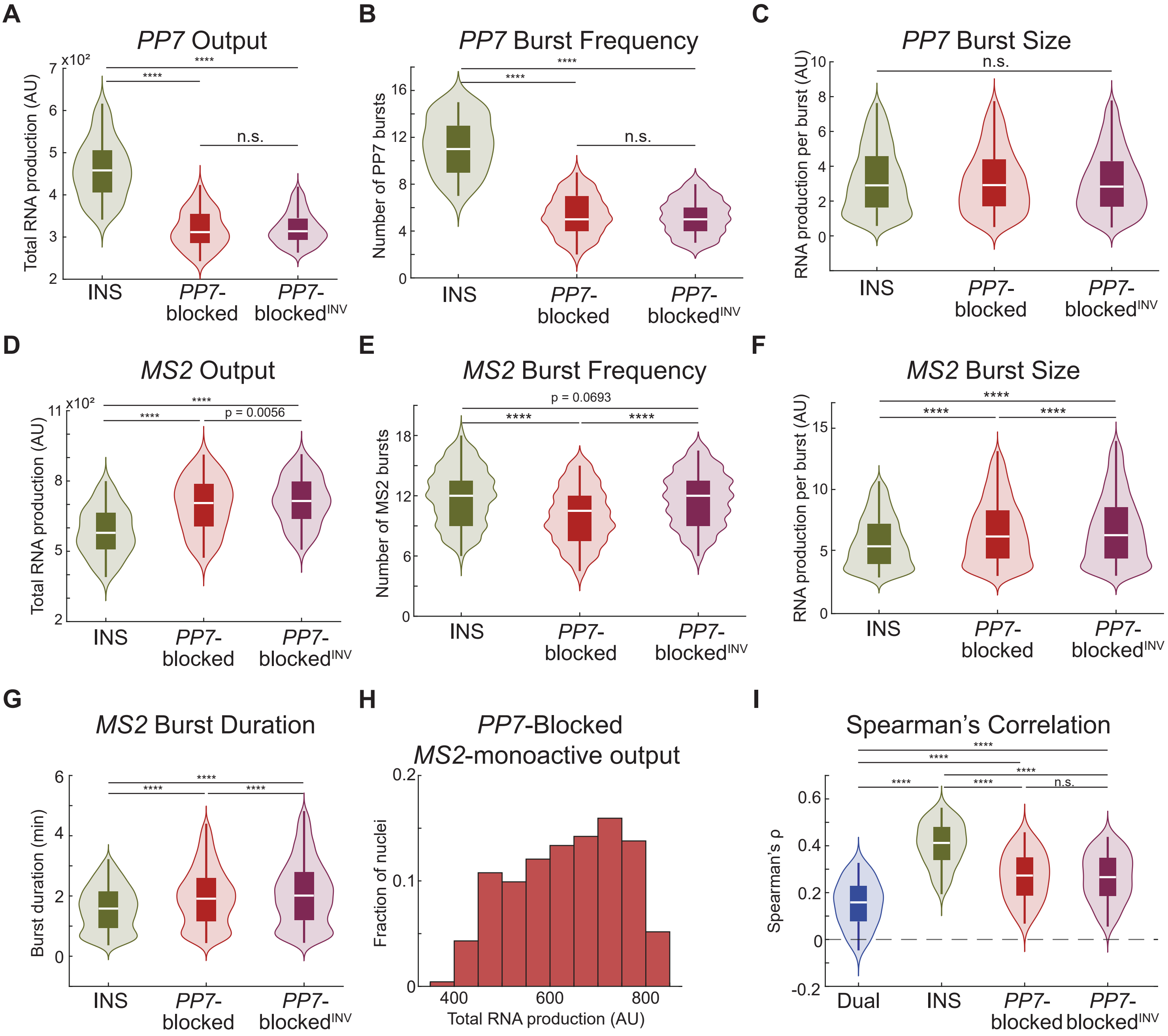


**Supplemental Figure 4.**
(A) Violin plot of total *PP7* output from individual *PP7*-active nuclei (n(INS) = 600, n(*PP7*-Blocked) = 355, n(*PP7*-Blocked$^{INV}$) = 549 nuclei).
(B) Violin plot of number of *PP7* bursts per nucleus. (n(INS) = 600, n(*PP7*-Blocked) = 355, n(*PP7*-Blocked$^{INV}$) = 549 nuclei).
(C) Violin plot of size of individual *PP7* bursting events. (n(INS) = 6889, n(*PP7*-Blocked) = 2516, n(*PP7*-Blocked$^{INV}$)= 3755 bursting events).
(D) Violin plot of total *MS2* output from *MS2*-active nuclei. (n(INS) = 575, n(*PP7*-Blocked) = 581, n(*PP7*-Blocked$^{INV}$)= 802 nuclei).
(E) Violin plot of number of *MS2* bursts per nucleus. (n(INS) = 575, n(*PP7*-Blocked) = 581, n(*PP7*-Blocked$^{INV}$)= 802 nuclei).
(F) Violin plot of size of individual *MS2* busting events.(n(INS) = 7106, n(*PP7*-Blocked) = 6245, n(*PP7*-Blocked$^{INV}$)= 9502 bursting events).
(G) Violin plot of length of individual *MS2* bursting events. (n(INS) = 7106, n(*PP7*-Blocked) = 6245, n(*PP7*-Blocked$^{INV}$)= 9502 bursting events).
(H) Histogram of total *MS2* output of *MS2*-monoactive nuclei from *PP7*-Blocked construct.
Hartigan's Dip Test of Unimodality indicated no significant deviation from unimodal distribution for either construct. D(*PP7*-Blocked) = 0.0169, p(*PP7*-Blocked) = 0.956, n(*PP7*-Blocked) = 232. $n_{boot}$ = 2000.
(I) Violin plot of Spearman's correlation coefficient of coactive nuclei. (n(Dual) = 361, n(INS) = 566, (*PP7*-Blocked) = 442, n(*PP7*-Blocked$^{INV}$) = 684 nuclei).
n.s. not significant, ****p<0.0001 by one-way ANOVA and Tukey-Kramer test.

### Supplemental figure 5

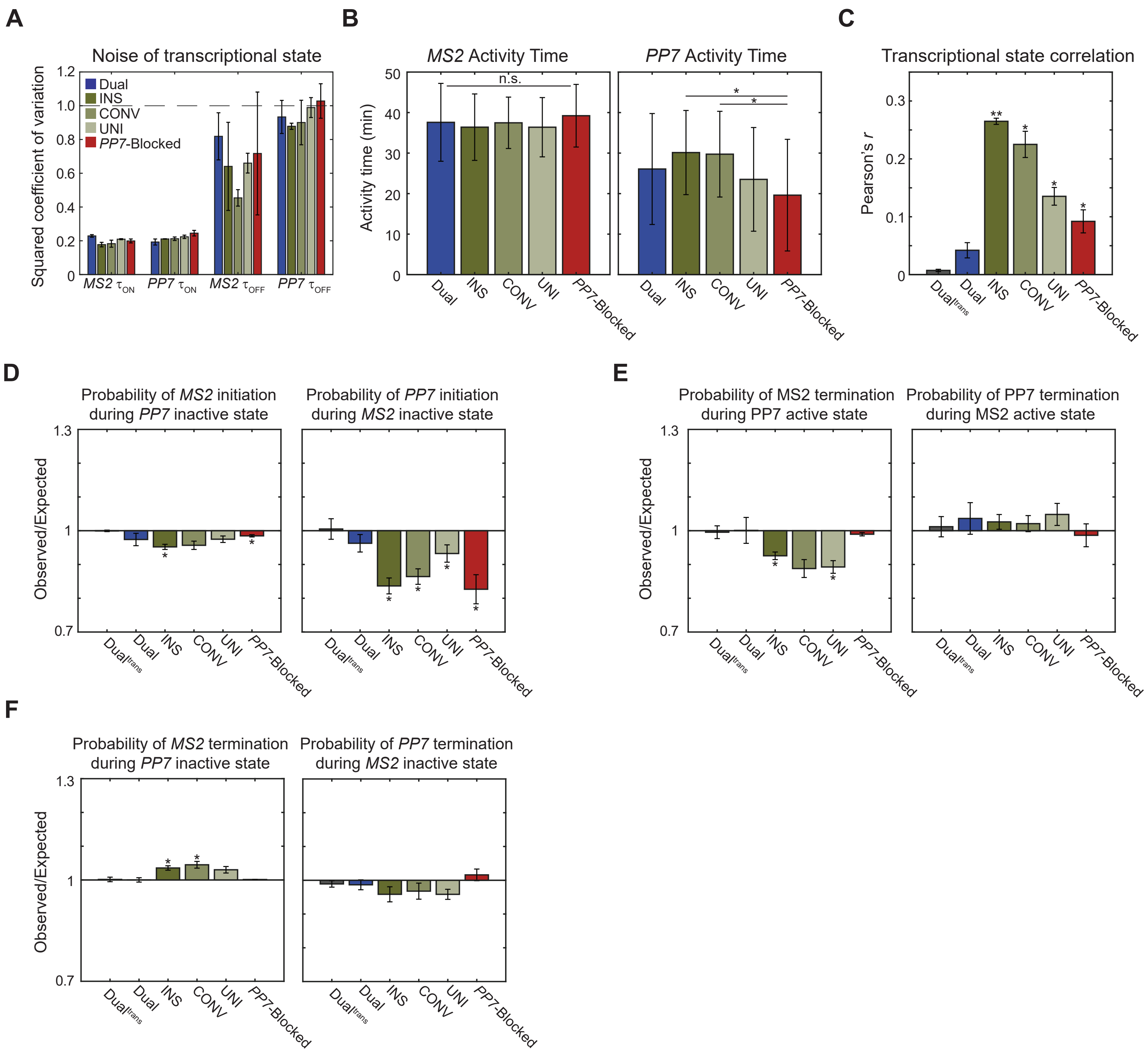


**Supplemental Figure 5.**
(A) Random varation measured as the squared coefficient of variation of the duration of each transcriptional state.
(B) Activity time of *MS2* (left) and *PP7* (right) for each construct.
(C) Pearson's correlation between *MS2* and *PP7* transcriptional state trajectories.
(D) Probability of (left) *MS2* promoter transitioning to ON state or (right) *PP7* promoter transitioning to ON state within time window of 27s while the other promoter is inactive.
(E) Probability of (left) *MS2* promoter transitioning to OFF state or (right) *PP7* promoter transitioning to OFF state within time window of 27s while the other promoter is active.
(F) Probability of (left) *MS2* transitioning to OFF state or (right) *PP7* promoter transitioning to OFF state within time window of 27s while the other promoter is inactive.
Probabilities are determined per biological replicate from minute 9 to 27 of nc 14.
Data are represented as mean ± SEM (N = 3 biological replicates). n.s. not significant, *p<0.05, **p<0.01 determined by one-way ANOVA and Tukey-Kramer test (B), one-sample t-test (C) and log-transformed, one-sample t-test (D,E,F).
